# Intramembrane protease SPP defines a cholesterol-regulated abundance control of the mevalonate pathway enzyme SQS

**DOI:** 10.1101/2021.07.19.452877

**Authors:** Dönem Avci, Ronny Heidasch, Martina Costa, Christian Lüchtenborg, Dipali Kale, Britta Brügger, Marius K. Lemberg

## Abstract

Intramembrane proteolysis regulates important processes such as signaling and transcriptional and posttranslational abundance control of proteins with key functions in metabolic pathways. This includes transcriptional control of mevalonate pathway genes, thereby ensuring balanced biosynthesis of cholesterol and other isoprenoids. Our work shows that, at high cholesterol levels, signal peptide peptidase (SPP) cleaves squalene synthase (SQS), an enzyme that defines the branching point for allocation of isoprenoids to the sterol and non-sterol arms of the mevalonate pathway. This intramembrane cleavage releases SQS from the membrane and targets it for proteasomal degradation. Regulation of this mechanism is achieved by the E3 ubiquitin ligase TRC8 that, in addition to ubiquitinating SQS in response to cholesterol levels, acts as an allosteric activator of SPP-catalyzed intramembrane cleavage of SQS. Cellular cholesterol levels increase in the absence of SPP activity. Hence, SPP-TRC8 mediated abundance control of SQS acts as a regulation step within the mevalonate pathway.

## Introduction

Cellular levels of key metabolites including lipids and sterols need to be tightly regulated in order to match the cellular demand. In particular, cholesterol, the most abundant individual lipid, has profound influence on membrane properties. Thus, cholesterol levels are modulated by uptake, endogenous synthesis, intracellular storage as cholesteryl ester, and efflux to apolipoproteins that altogether maintain membrane cholesterol homeostasis and prevent deleterious effects of hypercholesterolemia. Cholesterol is synthesized in the Endoplasmic Reticulum (ER) by the mevalonate pathway in a multistep process (1). One of the intermediates of cholesterol biosynthesis, farnesyl pyrophosphate (FPP), serves also as the building block of several additional molecules including dolichol and ubiquinone and it is utilized in post-translational modifications such as glycosylation and prenylation (2). Since these isoprenoids affect important biological activities, commitment to the non-sterol branch of the mevalonate pathway needs to be tightly controlled. On one side, enzymes of the mevalonate pathway are transcriptionally upregulated by the sterol regulatory element binding protein (SREBP) in response to low cholesterol (3). On the other hand, rate-limiting mevalonate pathway enzymes, HMG-CoA reductase (HMGCR) and squalene epoxidase (SQLE), are under post-translational feedback inhibition (4–7). When there is no demand for synthesis, the abundance of these key enzymes is downregulated by the ER-associated degradation (ERAD) pathway, a mechanism whereby proteins are targeted for turnover by the proteasome (8). While these homeostatic mechanisms determine the overall biosynthesis rate, SQLE has been suggested to influence the flux between the sterol- and non-sterol arm of the mevalonate pathway (5–7) (***Figure S1A***). However, a truncated SQLE form (SQLE-S) that persist cholesterol-induced degradation has been shown recently (9), posing the question of how pathway selection is solely accommodated at SQLE step. Underlying most of these homeostatic responses to fluctuating amounts of cholesterol is the sensing of its level by a number of regulatory ER membrane proteins via conserved sterol-sensing domains (SSD) (10). At low cholesterol level, SSD-mediated protein-protein interaction between Insig and SCAP is destabilized enabling trafficking of the SREBP from the ER to the Golgi, where its transcription factor domain is released and activated by regulated proteolysis by site-1 and site-2 proteases (11). Conversely, under conditions of high cholesterol, HMGCR is targeted to a redundant set of ERAD E3 ubiquitin ligases, leading to its extraction from the membrane and proteasomal turnover (7,12–15). In addition to these multiple ERAD dislocation mechanisms by the E3 ubiquitin ligases, cells have a non-canonical proteolytic arm of the ERAD pathway that is centered around signal peptide peptidase (SPP) (16). SPP is an aspartic intramembrane protease related to Alzheimer’s disease-associated γ-secretase (17). In addition to its role in processing signal peptides, SPP was shown to be also involved in protein degradation, functionally interacting with the ERAD E3 ubiquitin ligases TRC8 and MARCH6 (18–22). After these fundamental findings, further impact of SPP-triggered degradation on cellular physiology and the mechanism of its regulation remain as interesting research questions. In this study, we show that SPP and TRC8 trigger cholesterol-dependent proteasomal degradation of squalene synthase (SQS), an enzyme acting at the junction of sterol and non-sterol arms, thereby serving as a metabolic control at this branching point of the mevalonate pathway. We hypothesize that, this mechanism potentially contributes to pathway selection between the two branches, as it was also suggested for SQLE (4–7).

## Results

### Substrate proteomics identifies SQS as a physiological SPP substrate

To identify endogenous substrates for SPP, we compared the proteome of ER-derived microsomes from Hek293T cells, with or without treatment with the SPP inhibitor (Z-LL)_2_-ketone (17), using stable-isotope labeling by amino acids in cell culture (SILAC) and quantitative proteomics (***Figure 1A***). We identified 15 ER-resident proteins that showed an increased steady-state level of more than 20% in the inhibitor treated condition (see ***Supplementary file 1***). In addition to the previously identified SPP substrates heme oxygenase 1 (HO1) and syntaxin-18 (19,22,23), inhibition of SPP led to an increased level of SQS (***Figure 1B***), an ER-resident tail-anchored protein that defines the critical branching point of the sterol arm of the mevalonate pathway, by converting FPP into squalene (***Figure S1A***) (1). In contrast, 69 additional tail-anchored proteins identified were not significantly changed in their abundance (***Supplementary file 1***), indicating that SPP is highly specific.

**Figure 1.**
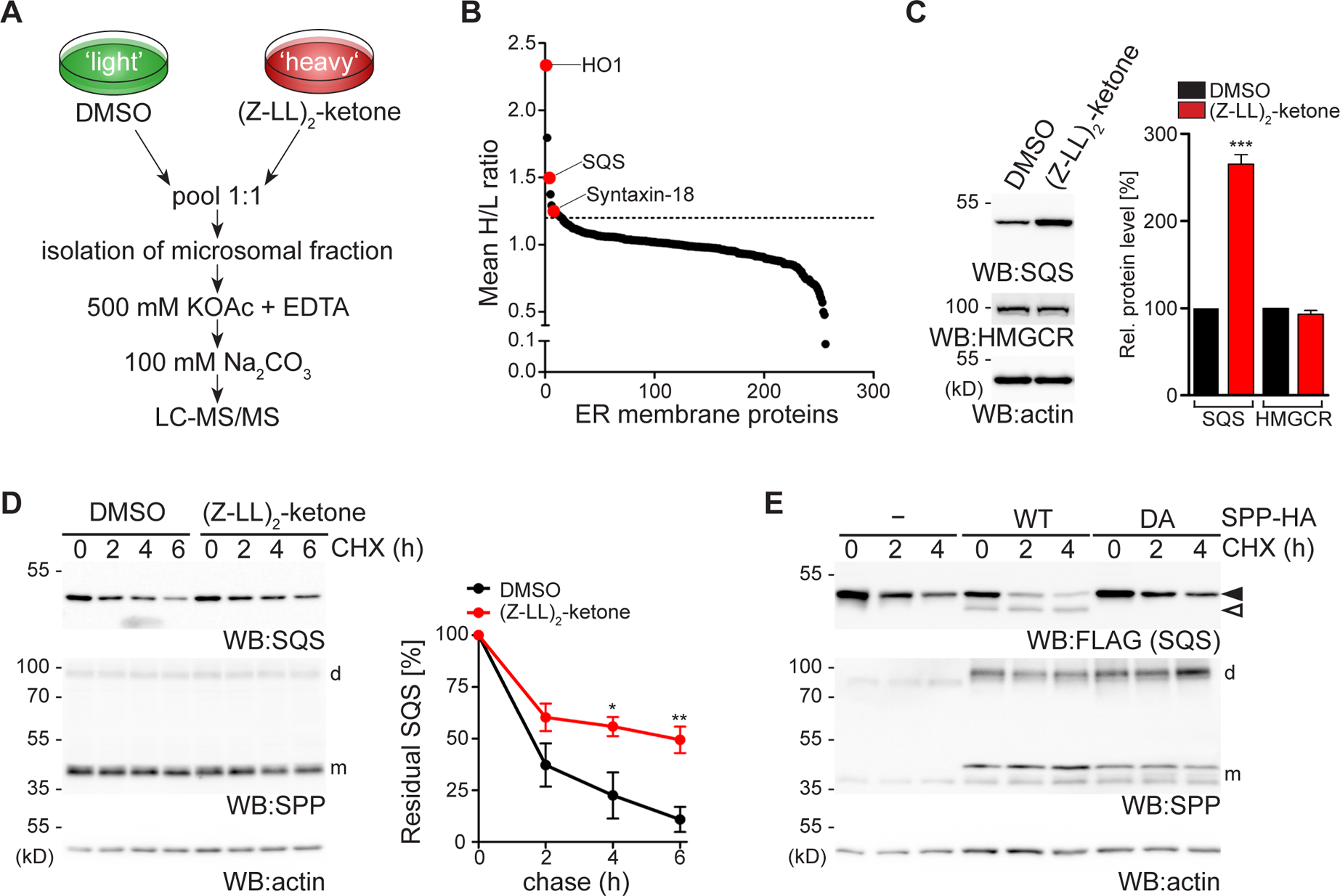
Differential organelle proteomics identifies SQS as an SPP substrate. **(A)** Experimental outline of the SILAC-based organelle proteomics. **(B)** ER-resident membrane proteins quantified by organelle proteomics (mean H/L ratio, *n*=2). **(C)** Western blot (WB) showing steady state analysis of endogenous SQS and HMGCR in Hek293T cells upon 16 h treatment with (Z-LL)_2_-ketone or DMSO. Actin was used as loading control. Quantification shows Mean ± SEM, *n*=4. **(D)** Cycloheximide (CHX) chase of endogenous SQS in Hek293T cells in presence of (Z-LL)_2_-ketone or vehicle control (DMSO). Monomeric and SDS-stable dimeric forms of SPP are indicated (m and d, respectively). Actin was used as loading control. Mean ± SEM, *n=3*. **(E)** CHX chase of ectopically expressed FLAG-tagged SQS in Hek293T cells co-expressing HA-tagged wild-type (wt) SPP and catalytic mutant SPP^D265A^ (DA) revealing processing of the full-length form (black arrow) to an N-terminal fragment (white arrow). See also figure S1.

Consistent with an SPP-mediated degradation of SQS, western blot analysis of (Z-LL)_2_-ketone treatment showed a significant increase of its steady-state levels, whereas HMGCR levels remained unchanged (***Figure 1C***). Consistent increase of SQS levels was observed also in HeLa and U2OS cells (***Figure S1B***). Furthermore, inhibition of SPP delayed SQS turnover in a cycloheximide chase assay (***Figure 1D***). In agreement with this, co-expression of a FLAG-tagged SQS construct with SPP destabilized the full-length form and a faster migrating band was observed, whereas ectopic expression of the catalytically inactive mutant SPP^D265A^ had no effect (***Figure 1E***). The tail-anchored protein Ubc6 was completely resistant to SPP-catalyzed cleavage (***Figure S1C***), supporting the notion that SPP is highly specific.

A key determinant of SPP substrates are helix-destabilizing residues in the N-terminal portion of the transmembrane (TM) domain, as well as specific amino acid residues surrounding the cleavage site (24). Mutation of either of two conserved TM serine residues (***Figure S1D***) matching this substrate consensus completely stabilized SQS in the ER (***Figure S1E, F***), indicating that a membrane-integral degradation signal (degron) determines its recognition by SPP. In order to confirm such a direct substrate-enzyme interaction, we performed co-immunoprecipitation experiments with the dominant-negative SPP^D265A^ mutant (18,23), revealing efficient trapping of endogenous SQS, whereas the ER protein CLIMP63 did not interact (***Figure S1G***). Taken together, these results show that SQS is a substrate for the SPP-dependent non-canonical ERAD pathway.

### SQS is targeted for degradation by Hrd1 as well the SPP-TRC8-dependent ERAD pathway

Previous studies showed that membrane insertion of SQS depends on the ER membrane protein complex (EMC) and defective insertion upon EMC loss leads to decreased stability of SQS, which was suggested to be independent of canonical ERAD pathways at this condition (25,26). In order to investigate how SQS turnover relates to the SPP-specific ERAD machinery, we generated SPP knockout Hek293T cells. Although non-conditional SPP knockout mice die after embryonic day 13.5 (27), SPP-deficient tissue culture cells grow normal, while accumulating the unprocessed substrate HO1 and show a defect in processing of the hepatitis C virus core protein (***Figure S2A,B***) (19,27). Consistent with our proteomics analysis, we observed significant steady-state increase of endogenous SQS, but not HMGCR in Hek293T SPP knockout cells (***Figure S2C***), whereas SQS mRNA level was not affected (***Figure S2D***). Although there was still substantial turnover of SQS in SPP knockout cells, cycloheximide chase assay showed a significant increase of the half-life compared to the wild type cells (***Figure 2A***). This result indicates that, SQS is subject to more than one ERAD pathway as had been observed for HMGCR (***Figure S1A***). We next generated knockout cells of both E3 ubiquitin ligases that had been linked to SPP, namely TRC8 and MARCH6 (18,20,21), and analyzed SQS protein stability and mRNA expression levels (***Figure 2B and Figure S2E***). Similar to SPP knockout, we observed a partial stabilization of SQS in ΔTRC8 cells, whereas MARCH6 knockout did not show any significant effect (***Figure 2B and Figure S2F***). Combined knockout of TRC8 and MARCH6 did not further stabilize SQS (***Figure 2B***). On the other hand, knockdown of the Hrd1 E3 ubiquitin ligase by siRNA transfection also interfered with SQS degradation (***Figure S2G***). Strikingly, Hrd1 knockdown in the TRC8 and MARCH6 double-deficient cells almost completely stabilized SQS (***Figure 2C***). Consistent with SPP and TRC8 acting in one linear ERAD pathway, addition of the SPP inhibitor (Z-LL)_2_-ketone did not show any effect in 1′SPP or 1′TRC8 cells but stabilized SQS in ΔMARCH6 cells (***Figure S2H-J***). Taken together, these results show that SQS is subject to at least two independent branches of the ERAD pathway, the Hrd1-dependent dislocation route and the SPP-TRC8 pathway (18,19).

**Figure 2.**
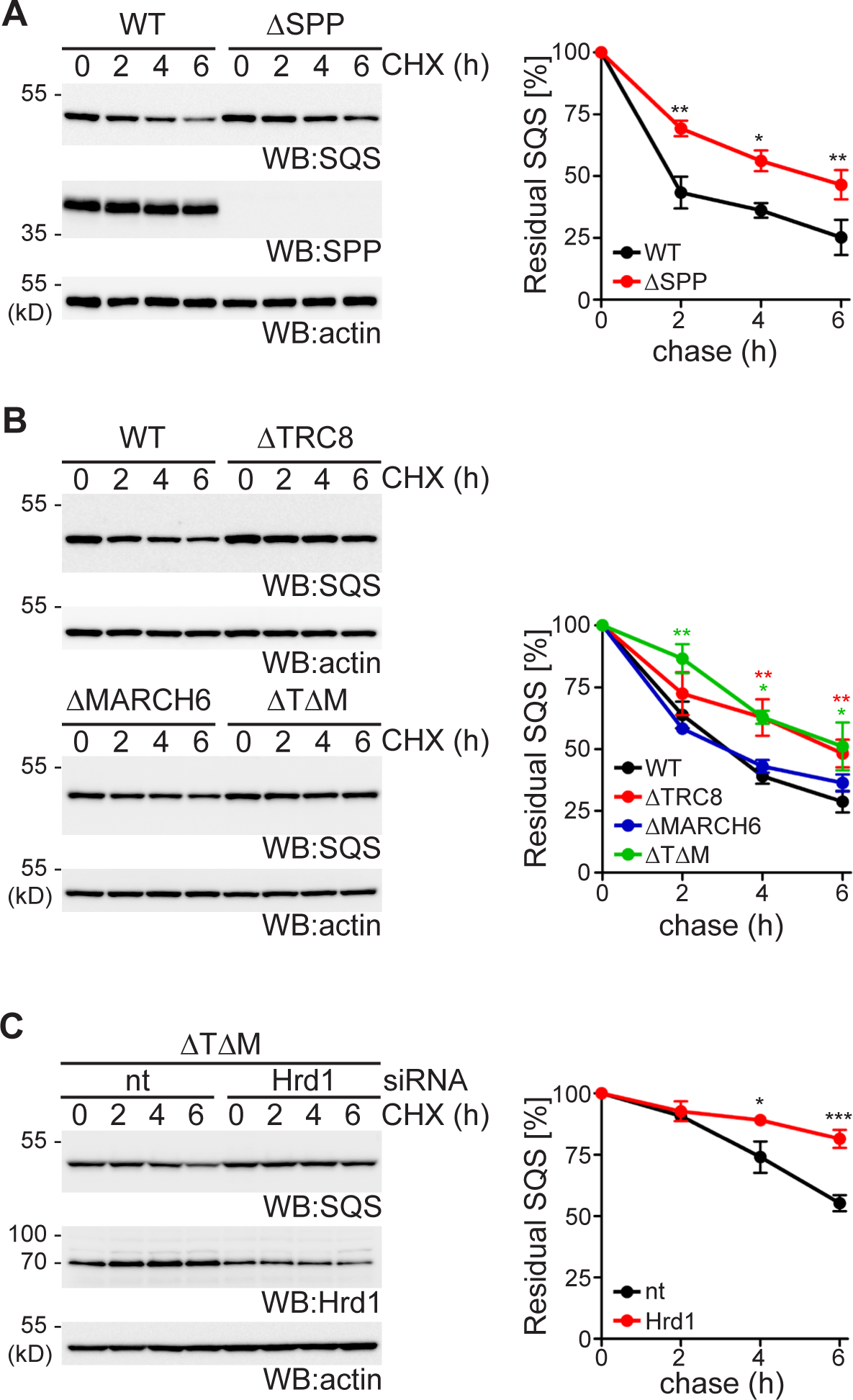
SPP and TRC8 define a non-canonical ERAD pathway that acts on SQS in parallel to Hrd1 pathway. **(A)** Stability of endogenous SQS in Hek293T wt and Hek293T1′SPP cells assessed by cycloheximide (CHX) chase assay. Actin was used as loading control. Mean ± SEM, *n=3*. **(B)** SQS degradation assay in Hek293T wt, 1′TRC8, 1′MARCH6, and 1′TRC81′MARCH6 (1′T1′M) double-deficient Hek293T cells. Mean ± SEM, *n=3.* **(C)** SQS degradation assay in 1′T1′M cells transfected with non-targeting (nt) or Hrd1 siRNA. Mean ± SEM, *n=3.* See also figure S2.

### Cholesterol activates SPP-TRC8 activity

To assess whether SPP-TRC8-mediated SQS turnover is regulated, we studied its degradation rate in response to cholesterol levels. For this, we used lipoprotein-free medium and conditions blocking endogenous cholesterol synthesis, while synthesis of essential non-sterol isoprenoids is permitted and no toxicity was observed (***Figure 3A and Figure S3A***) (28,29). SQS was significantly more stable in Hek293T cells grown in cholesterol-depletion medium when compared to cholesterol replete conditions (***Figure 3B***). The effect observed was comparable to cholesterol-induced ERAD of full-length SQLE (***Figure 3B***) (7), whereas the truncated SQLE-S form was stable, as observed before (9). In contrast, in SPP- or TRC8-deficient cells, SQS turnover was delayed and completely uncoupled from cholesterol levels, whereas cholesterol-induced ERAD of SQLE was unaffected by both knockouts (***Figure 3C, D***). This effect was specific to cholesterol, since 25-hydroxycholesterol, which serves as a potent trigger of HMGCR turnover (30), did not affect SQS stability (***Figure S3B***), as previously had been observed for SQLE (5). In Hrd1 deficient cells, on the other hand, cholesterol addition still led to accelerated degradation of SQS, similar to what we observed in wild type cells (***Figure 3E***). Taken together, our results show that for both enzymes a basal turnover exists. At high cholesterol conditions, SQLE is degraded by a MARCH6- and p97-dependent dislocation route (7), and SQS becomes subject to SPP-TRC8-triggered turnover. Consistent with a cholesterol-induced ubiquitination of SQS, we observed increased levels of several higher molecular weight SQS species in a ubiquitin-specific pulldown in wild type cells (***Figure 3F***). This difference was diminished in ΔTRC8 cells but still observed in the absence of SPP (***Figure 3F***). These results suggest that, TRC8-catalyzed ubiquitination serves as the sterol-sensitive step that is succeeded by SPP-catalyzed release from the ER membrane.

**Figure 3.**
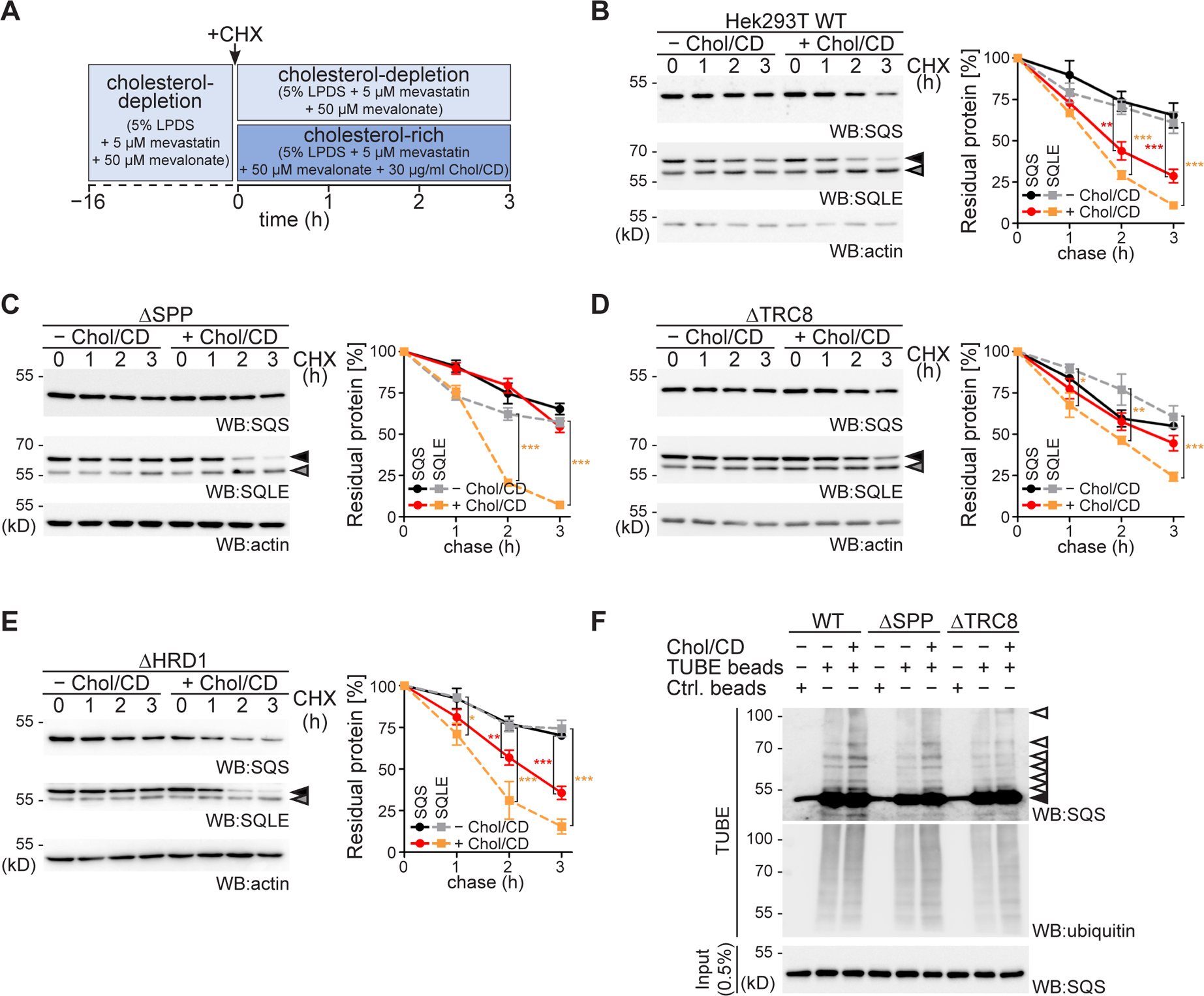
Cholesterol activates SPP-TRC8 complex for regulatory degradation of SQS. **(A)** Experimental outline of the sterol starvation and cholesterol repletion assay. LPDS, lipoprotein-depleted serum; Chol/CD, cholesterol complexed to methyl-β-cyclodextrin. **(B)** Degradation of endogenous SQS and the full-length form of SQLE (black arrow) were assayed in Hek293T wt cells treated as outlined in (**A**) and analyzed by western blot (WB) analysis. Actin was used as loading control. Grey arrow, cholesterol-insensitive truncated SQLE-S. Mean ± SEM, *n=3*. **(C)** SQS and SQLE degradation assay in Hek293TΔSPP cells as in (**B**). **(D)** SQS and SQLE degradation assay in Hek293TΔTRC8 cells as in (**B**). **(E)** SQS and SQLE degradation assay in Hek293T1′HRD1 cells as in (**B**). **(F)** Hek293T wt, ΔSPP, and ΔTRC8 cells were grown in sterol-depletion medium and treated with epoxomicin and incubated in cholesterol depletion or repletion as outlined in (**A**) before cell lysates were subjected to affinity enrichment using TUBE2 agarose beads revealing ubiquitinated species (white arrows). Control (Ctrl.) agarose beads were used to confirm TUBE2 specificity. See also figure S3.

### TRC8 triggers SPP-catalyzed SQS cleavage

Previous studies showed that TRC8 levels are modulated by sterols and it interacts with sterol biogenesis regulating proteins Insig, SREBP2 and SCAP (31,32). It was also shown to affect the processing and stability of SREBP precursor proteins and was suggested to fine tune SREBP response upon prolonged sterol depletion (31,32). Therefore, we analyzed the potential sensor role of TRC8 on sterol regulated degradation of SQS. Expression of TRC8 harboring a tyrosine-32-glutamate (Y32E) mutation, which previously has been suggested to compromise its SSD (32), showed significantly reduced activity in triggering SQS degradation in ΔTRC8 cells compared to the wild-type construct (***Figure 4A, B***).

**Figure 4.**
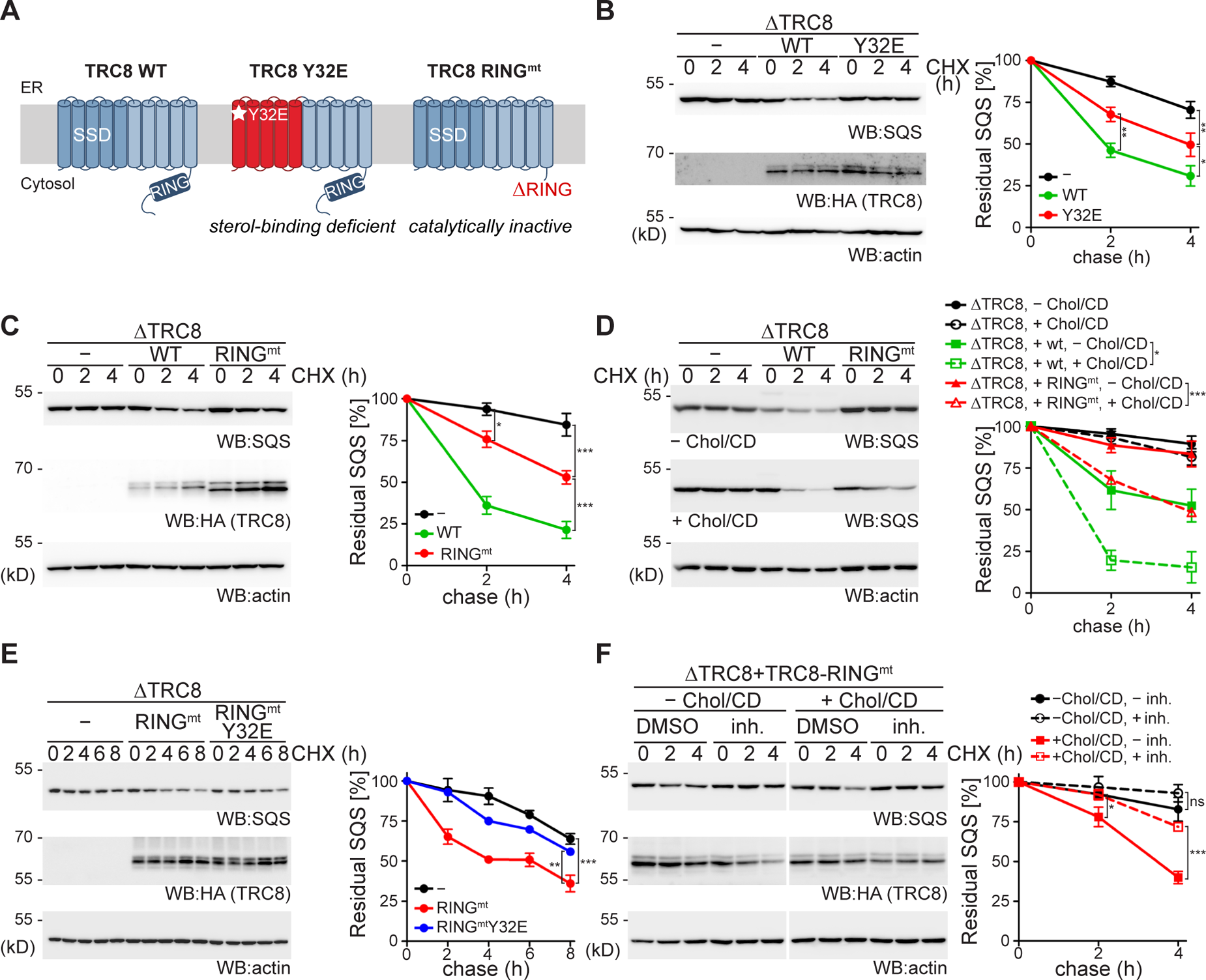
TRC8 acts as an allosteric activator of SPP-catalyzed SQS cleavage. **(A)** Outline of TRC8 constructs used. **(B)** Doxycycline-induced expression of HA-tagged TRC8 wt or the Y32E mutant in Hek2931′TRC8 cells. Cells were subjected to CHX chase and assayed for endogenous SQS. Mean ± SEM, *n=3*. **(C)** Doxycycline-induced expression of HA-tagged TRC8 wt or RING domain mutant (RING^mt^) in Hek2931′TRC8 cells. Cells were subjected to CHX chase and assayed for endogenous SQS. Mean ± SEM, *n=3*. **(D)** Doxycycline-induced expression of HA-tagged TRC8 wt or the RING^mt^ in Hek2931′TRC8 cells in cholesterol depletion or repletion. Cells were assayed for endogenous SQS by CHX chase as outlined in Figure 3A. Mean ± SEM, *n=3*. **(E)** Doxycycline-induced expression of HA-tagged TRC8 RING^mt^ or the RING^mt^ Y32E double mutant in Hek2931′TRC8 cells. Cells were subjected to CHX chase assay and assayed for endogenous SQS. Mean ± SEM, *n=3*. **(F)** Doxycycline-induced expression of HA-tagged TRC8-RING^mt^ in Hek2931′TRC8 cells was assayed for endogenous SQS by CHX chase in presence or absence of (Z-LL)_2_-ketone (inh.). Mean ± SEM, *n=4*.

Interestingly, deletion of TRC8’s catalytic RING domain (RING^mt^) only partially abrogated rescue of the knockout phenotype (***Figure 4A, C***), indicating that TRC8 has also a non-catalytic function in SQS turnover by sensing the cholesterol level. Consistent with this hypothesis the non-catalytic function of TRC8-RING^mt^ was entirely cholesterol dependent (***Figure 4D***) and the RING^mt^-Y32E double mutant of TRC8 did not rescue SQS degradation in ΔTRC8 cells (***Figure 4E***). In addition, SPP inhibition under high cholesterol conditions in ΔTRC8 cells expressing TRC8-RING^mt^ led to a significant stabilization of SQS level (***Figure 4F***), suggesting an allosteric activation of SPP proteolytic activity mediated through TRC8. Altogether, these results indicate that high cholesterol conditions enhance the proteolytic processing of SQS by SPP. Moreover, for complete SQS turnover, both the E3 ubiquitin ligase and the sterol sensing-activity of TRC8 are required.

### SPP-mediated SQS degradation regulates the mevalonate pathway

Next, we tested the influence of SPP mediated SQS abundance control on cellular cholesterol levels by analyzing the lipidome of wild type and ΔSPP Hek293T cells (***Figure S4A-E***). In order to uncouple cellular cholesterol level from receptor-mediated uptake (1), cells were grown in lipid-depleted medium. Total cholesterol levels were significantly increased in all three ΔSPP clones (***Figure S4F***). The difference was even more pronounced in the lipidomic analysis upon subcellular fractionation, showing a striking increase of cholesterol in both endomembrane and plasma membrane fractions (***Figure 5A***). The levels of cholesterol in the cell needs to be tightly controlled and even small changes can have profound effects. As a way to test how ablation of SPP affects plasma membrane properties, we measured uptake of fluorescently labelled dextran from the medium. Consistent with cholesterol-dependent endocytosis (33), in Hek293T cells grown in rich medium 70-kDa dextran was visible in various vesicles, whereas no internalization was observed in cholesterol-depleted medium (***Figure 5B***). Strikingly, significantly more 70-kDa dextran was internalized in ΔSPP cells, compared to wild type, in cholesterol depletion conditions (***Figure 5B***), indicating that increased plasma membrane cholesterol level, due to uncoupled cholesterol biosynthesis in ΔSPP cells, facilitates endocytosis under these conditions. This increased level of cholesterol-dependent endocytosis in ΔSPP cells was even more pronounced for the uptake of 10-kDa dextran (***Figure 5B***), which, in addition to macropinocytosis, also becomes subject to a less-well defined pinocytosis route (33). This shows that, SPP-mediated SQS abundance control is important to fine-tune cellular cholesterol levels with a broad impact on membrane properties. Total lipidome analysis of Hek293T wt and ΔSPP cells did not show any major differences in glycerophospholipid and storage lipid content. However, we observed an overall reduction in sphingolipid levels in ΔSPP cells (***Figure S4G***), as well as an increase in long chain phosphatidylserine (36:1, see ***Figure S4C***) that has been linked to positioning of cholesterol in the plasma membrane (34). The mechanism of how SPP influences sphingolipid homeostasis and coupling of cholesterol to membrane composition remain important questions for future studies.

**Figure 5.**
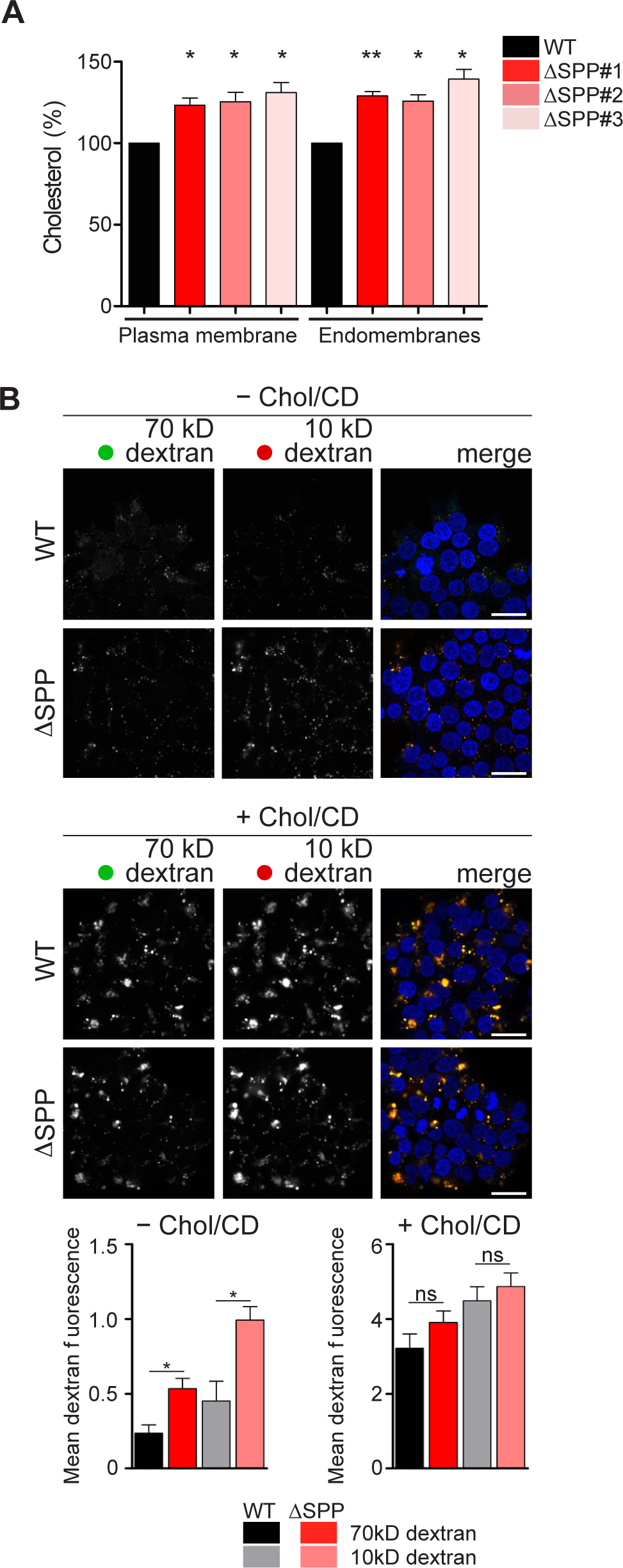
SPP-mediated SQS quantity regulation plays a crucial role in the control of cholesterol homeostasis. **(A)** Lipidomic analysis of subcellular fractionation samples comparing Hek293T wt and three ΔSPP cell lines (clone #1, #2, #3). Mean ± SEM, *n*=*3*. **(B)** Hek293T wt and ΔSPP cells were grown in sterol-depletion medium (top panel) or in cholesterol-rich medium (bottom panel) and incubated with fluorescently labelled 70-kDa dextran (green) and 10-kDa dextran (red). Intracellular dextran was quantified after 30 min uptake. Nuclear DNA was stained by Hoechst 33342 (blue). Mean ± SEM, *n* = 10 fields of view. Scale bar, 100 µm. See also figure S4.

## Discussion

Regulation of metabolite levels according to cellular needs is crucial to maintain homeostasis. Here we show that, the regulatory arm of the ERAD pathway (ERAD-R) centered around SPP (16) acts as a so far unrecognized metabolic regulation in the mevalonate pathway (***Figure 6***). Since the SQS-catalyzed conversion of two molecules of FPP to squalene determines to which extent the mevalonate pathway generates sterol or other biologically important non-sterol molecules including dolichol and ubiquinone, we hypothesize that it potentially serves as a metabolic switch at the branching point of the mevalonate pathway (***Figure 6***). This regulated degradation can act as a fast-reacting fine-tuning process to control the flux of the FPP intermediate into two arms of the mevalonate pathway according to the cellular needs. While HMGCR, as the first rate-limiting enzyme of the multistep pathway, determines the overall flux of acetyl-CoA into isoprenoid biosynthesis, regulated turnover of SQS defines a control step that shunts FPP between the sterol and non-sterol branches. Cholesterol levels are increased in SPP knockout cells, due to increased SQS abundance. This shows that, in addition to the control of the rate-limiting enzyme HMGCR by multiple ERAD E3 ubiquitin ligases (7,12–14), cholesterol also acts as a negative feedback regulator on the branching-point enzyme SQS. Consistent with this, we observed that the ERAD E3 ubiquitin ligases TRC8 and Hrd1 show an overlapping substrate spectrum, potentially allowing integration of different signals to control the abundance of SQS and ensuring plasticity of the response. The change in cholesterol levels in SPP knockout cells is also reflected to an increased capacity of cholesterol-dependent endocytosis compared to wild type cells, proving the physiological relevance of this metabolic regulation. How the complex cellular metabolism of non-sterol mevalonate products influences this control and whether there are compensating mechanisms that suppress an SPP knockout phenotype, as well as how an SPP upregulation observed in cancer cells (35,36) impacts this mechanism, remain to be important questions for future research.

**Figure 6.**
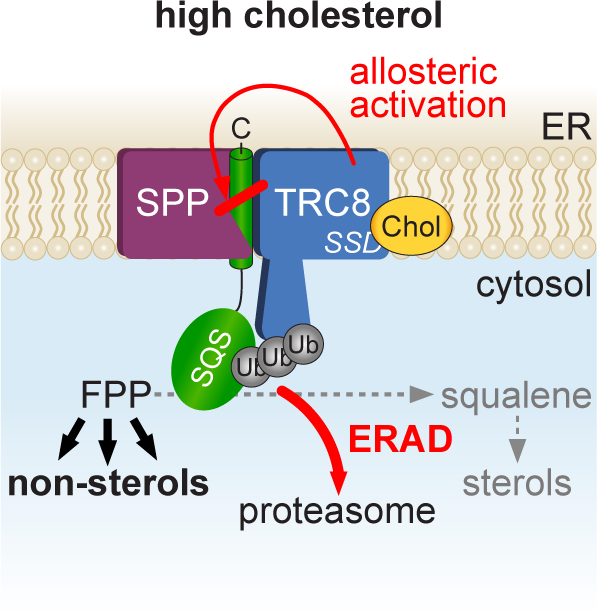
Model for SPP-triggered cholesterol dependent degradation of SQS. At high levels, cholesterol is sensed by the sterol-sensing domain (SSD) of TRC8 in Hek293T wt cells leading to ubiquitination and subsequent SPP-mediated ERAD-R of SQS.

We reveal that SPP is activated partially by a non-catalytic sterol-sensing function of TRC8 to cleave SQS. This observation answers a long-standing question about the physiological relevance of the predicted SSD in TRC8 (32). As noted previously, TRC8 levels are also regulated by sterol amounts in the cell. We believe TRC8 mediated degradation of SQS is part of a complex multi-level metabolic regulation, acting as distinct control steps during the time course of cellular response to sterol depletion and repletion. Different to ERAD of HMGCR (4) and activation of the transcription factor SREBP (11), 25-hydroxycholesterol does not show an effect on the SPP-TRC8-mediated protein abundance control. This difference indicates that, for this ERAD-R mechanism the end product of cholesterol biosynthesis is sensed and not the general flux through the mevalonate pathway is monitored. The molecular mechanism of how TRC8’s SSD activates ERAD-R remains to be determined. However, we assume that recognition of single-pass TM substrates by the SPP-TRC8 complex is independent of SCAP/Insig adaptor proteins that are known to control HMGCR and SREPB (37). Since SPP specificity is determined by TM substrate features (24), we suggest that the cholesterol-induced increase of the SQS processing rate is caused by allosteric activation of SPP by the membrane-integral SSD of TRC8. Presenilin, a mechanistically related aspartic intramembrane protease serving as the active subunit of the γ-secretase complex, is also modulated by cholesterol, which impacts generation of neurotoxic amyloid-β peptides in Alzheimer’s disease (38). A recent γ-secretase structure revealed three distinct cholesterol-binding sites in the non-catalytic subunit Aph-1 adjacent to presenilin (39), suggesting that allosteric regulation of aspartic intramembrane proteases by lipids is a more general principle. Over all, intramembrane proteolysis emerges as a fast and irreversible post-translational modification that controls the activity and abundance of various organellar membrane proteins (16).

Human cancer association studies showed that SPP expression is increased in glioblastoma, lung, and breast cancer cells (35,36). Based on our findings, we hypothesize that reduced SQS level upon increased SPP expression and concomitant increased flux of FPP into dolichol and ubiquinone synthesis may help to handle the increased metabolic load of rapidly growing cancer cells. Consistent with this idea, a recent study showed that synthesis through the mevalonate pathway is shifted towards ubiquinone to sustain viability of p53-deficient cancer cells exposed to metabolic stress (40). Likewise, heme metabolism may contribute to settle the microenvironment for cancer cell growth. These new insights in the role of SPP in cellular metabolism control offer the prospect of novel therapeutic intervention strategies.

## Materials and Methods

### Antibodies

The following antibodies were used in this study: rabbit polyclonal anti-SPP (kind gift of C. Schaller), mouse monoclonal anti-FLAG (M2, Sigma-Aldrich, #F1804), rat monoclonal anti-HA (3F10, Sigma-Aldrich, #11867423001), mouse monoclonal anti-CLIMP63 (G1/296, Enzo Life Sciences, #ENZ-ABS669-0100), mouse monoclonal anti-β actin (AC-15, Sigma-Aldrich, #A1978), mouse monoclonal anti-ubiquitin (P4D1, Santa Cruz Biotechnology, #sc-8017), rabbit polyclonal anti-HO1 (Enzo Life Sciences, #ADI-SPA-896-F), rabbit polyclonal anti-SQLE (Proteintech, #12544-1-AP), rabbit monoclonal anti-SQS (Abcam, #ab109723), rabbit monoclonal anti-SQS (Abcam, #ab195046), mouse monoclonal anti-GFP (Sigma-Aldrich, #11814460001), mouse monoclonal anti-HMGCR (Merck, #MABS1233), rabbit polyclonal anti-Synoviolin (Hrd1) (Bethyl Laboratories, #A302-946A), AlexaFluor488 goat anti-mouse IgG (H+L) (Invitrogen, #A-11029), mouse monoclonal S-peptide epitope antibody (Thermo Fischer # MA1-981).

### Plasmids

Plasmids based on pcDNA3.1 (Invitrogen) encoding human SPP with a triple HA-tag inserted between residue 373 and the C-terminal KKEK ER-retention signal (SPP-HA) was described previously (18). The active-site mutant SPP D265A was introduced by site-directed mutagenesis (Stratagene). Human SQS (*FDFT1*, Gene ID 2222, full-ORF Gateway cDNA clone 101218403) and Ubc6-FLAG (Gene ID 51465, IMAGE 8322482) with a N-terminal triple FLAG-tag were inserted as *EcoR*I/*Xho*I-fragment into pcDNA3.1. TM mutants of SQS were generated by site-directed mutagenesis. For the generation of doxycycline-inducible cell lines, we introduced constructs for human TRC8 (Gene ID 11236), TRC8-RING^mt^ lacking residues 547 to 580 (gifts from R. Gemill (41)), and the Y32E SSD mutant generated by site-directed mutagenesis (this study), with a C-terminal single HA-tag respectively, into pcDNA5/FRT/TO (Invitrogen). Plasmids encoding an ER-targeting signal sequence fused to an RFP followed by a KDEL ER-retention signal (RFP-KDEL) (42) and the N-terminal 195 residues of the hepatitis C virus polyprotein (Glasgow strain genotype 1a) corresponding to the core protein and 4 amino acids of the envelope protein E1 fused to an N-terminal triple FLAG-tag (43) have been described before.

### Cell culture and transfection

Hek293T and Hek293 Flp-In T-REx cells were maintained without antibiotics in Dulbecco’s modified Eagle’s medium (DMEM) (Invitrogen) supplemented with 10% fetal bovine serum (FBS) (Invitrogen) and grown as monolayers at 37°C and 5% CO_2_. Hek293 Flp-In T-REx cells were additionally supplemented with 100 µg/ml hygromycin B (Invitrogen) and 10 µg/ml blasticidine (Gibco). Transient transfections were performed using 40-kDa linear polyethyleneimine (PEI MAX, Polysciences) (44). Typically, 1 µg encoding substrate and 500 ng of plasmid encoding wt or mutant SPP were used per well of a 6-well plate. Total DNA (2 µg/well) was held constant by adding empty plasmid. If not stated otherwise, cells were harvested 24 h after transfection. For inhibition of the proteasome and SPP, 2 µM epoxomicin and 50 µM (Z-LL)_2_-ketone (Merck Millipore) were added from stock solutions in dimethyl sulfoxide (DMSO). As vehicle control, the same amount of DMSO was used. For the induction of HO1, 20 µM hemin (ferriprotoporphyrin IX chloride, Sigma-Aldrich) was added to the cells from a 20 mM stock solution in 0.05 M NaOH and incubated overnight. For immunofluorescence staining, Hek293T cells were seeded onto 12 mm cover slips in 24-well plates and transfected with 500 ng plasmid encoding substrate and 150 ng RFP-KDEL. Total amount of DNA (1 µg/well) was held constant with empty plasmid.

### CRISPR/Cas9-mediated gene editing

CRISPR sgRNA oligonucleotide sequences targeting SPP (*HM13* gene), TRC8, or MARCH6 were designed using the E-CRISP online tool (45). The sgRNAs were cloned into the *Bbs*I-digested px459 (pSpCas9(BB))-2A-Puro V2.0, (Addgene #62988) vector, verified by sequencing and used for transient transfection of Hek293T cells and Hek293 Flp-In T-REx cells. sgRNA-targeted cell populations were selected with 3 µg/ml puromycin and clonal cell lines were generated by limited dilution. Knockout clones for SPP were identified by immunoblotting and validated by sequencing of genomic DNA. Validation of TRC8 and MARCH6 knockout cells was based on indel detection. Identification of indels was performed using the online tools TIDE (46), CRISP-ID (47), and ICE v2 (48). The following sgRNAs were used: SPP-1 (clone #1 and #3) GCCCTCAGCGATCCGCATAA, SPP-2 (clone #2) GCCTGAAACAATCACCAGCC, TRC8 CCAGACATACTACGAGTCTT, MARCH6 TATCATCCTTGTGTATGTAC. The Hrd1 knockout cells were described previously (49).

### Generation of stable Hek293 Flp-In T-REx cell lines

For the generation of doxycycline-inducible stable cell lines, Flp-In T-REx Hek293ΔTRC8 cells were co-transfected with pOG44 (Invitrogen) and either pcDNA5/FRT/TO/TRC8, pcDNA5/FRT/TO/TRC8-RING^mt^, or pcDNA5/FRT/TO/TRC8-Y32E and cells were selected with hygromycin B (100 µg/ml, Invitrogen).

### siRNA knockdown of Hrd1

For the knockdown of Hrd1 (*SYVN1* gene), cells were transfected with an ON-TARGETplus siRNA smart pool (L-007090-00-0005) obtained from Dharmacon. As control, non-targeting siRNA pool (D-001810-01) was used. Cells were seeded onto poly-L-lysine (PLL)-coated 12-well plates in DMEM supplemented with 10% FBS and transfected using the RNAiMAX reagent (Life Technologies) according to manufacturer’s instructions. After 24 h, medium was exchanged to fresh medium. Cells were harvested 48 h after transfection. The final molarity of siRNA per well was 20 picomol.

### Mass spectrometry-based substrate screen

For the identification of SPP substrates, we used stable isotope labelling by amino acids in cell culture (SILAC)-based quantitative organelle proteomics. For this, Hek293T cells were grown for five doublings in medium supplemented with either heavy amino acids (^13^C_6_^15^N_4_-L-Arg and ^13^C_6_^15^N_2_-L-Lys; Silantes) or unlabelled amino acids before applying (Z-LL)_2_-ketone (50 µM) or DMSO as vehicle control. For each condition ((Z-LL)_2_-ketone or DMSO), 3 x 150 mm plates were used.

If not stated otherwise, all steps were performed at 4°C. At day of harvest, cells were washed with ice-cold 1x PBS (10 mM NaH_2_PO_4_, 1.4 mM KH_2_PO_4_, 2.7 mM KCl, 140 mM NaCl, pH 7.4), harvested in PBS-EDTA (1x PBS, 1 mM EDTA, pH 8.0, 0.2 g/l glucose) and an equal number of cells were resuspended in ice-cold hypotonic buffer (10 mM HEPES-KOH, pH 7.4, 1.5 mM MgCl_2_, 10 mM KOAc, 0.5 mM dithiothreitol (DTT)) containing 10 µg/ml phenylmethylsulfonyl fluoride (PMSF) and Complete EDTA-free protease cocktail (Roche). Cells were incubated on ice for 10 min and lysed by passing five times through a 27-Gauge needle, followed by centrifugation for 10 min at 400 x *g* to remove cell debris and nuclei. The supernatant was laid on top of a low salt sucrose cushion (500 mM sucrose, 50 mM HEPES-KOH, pH 7.4, 50 mM KOAc, 5 mM Mg(OAc)_2_) and centrifuged for 20 min at 100,000 x *g* in a 70Ti rotor (Beckman Coulter). The resulting membrane pellet was resuspended in 1 ml rough microsome (RM) buffer (250 mM sucrose, 50 mM HEPES-KOH, pH 7.4, 50 mM KOAc, 2 mM Mg(OAc)_2_, 1 mM DTT) and extracted by 500 mM KOAc and 50 mM EDTA on ice for 15 min, followed by centrifugation through a high salt sucrose cushion (500 mM sucrose, 50 mM HEPES-KOH, pH 7.4, 500 mM KOAc, 5 mM Mg(OAc)_2_) for 20 min at 100,000 x *g*. The pellet thereof was resuspended in 1 ml freshly prepared, ice-cold sodium carbonate (100 mM) buffer and transferred onto an alkaline sucrose cushion (125 mM sucrose, 100 mM Na_2_CO_3_) followed by centrifugation for 20 min at 100,000 x *g*. The pellet was resuspended in RM buffer and snap frozen in liquid nitrogen.

For proteomics, pellets in RM buffer were diluted in 2x SDS-sample buffer (see below), separated by sodium dodecyl sulfate (SDS) polyacrylamide gel electrophoresis SDS-PAGE for 1 cm and samples were excised from the gel before reducing with DTT and alkylation with iodoacetamide. Digestion with trypsin was done overnight at 37°C, followed by quenching with 0.1% trifluoracetic acid (TFA; Biosolve, Valkenswward), and the supernatant was dried in a vacuum concentrator before LC-MS analysis. Nanoflow LC-MS^2^ analysis was performed using an Ultimate 3000 LC system coupled to an QExactive HF mass spectrometer (Thermo Scientific). Samples were dissolved in 0.1% TFA and loaded onto a C18 Acclaim PepMap100 trap-column (Thermo Fisher Scientific) with a flow rate of 30 µl/min 0.1% TFA. Subsequently, peptides were eluted and separated on an C18 Acclaim PepMap RSLC analytical column (75 µm x 250mm, Thermo Fisher Scientific) with a flow rate of 300 nl/min in a 120 min gradient of 3% buffer A (0.1% formic acid) to 40% buffer B (0.1% formic acid/acetonitrile). The mass spectrometer was operated in data-dependent acquisition mode, automatically switching between MS and MS^2^. Collision induced dissociation MS^2^ spectra were generated for up to 20 precursors with normalized collision energy of 29 %. Processing of RAW files was performed using MaxQuant (v. 1.5.3.30). MS^2^ spectra were searched against the Uniprot human proteome database and the contaminants database by Andromeda search engine with the following parameters: Carbamidomethylation of cysteine residues, acetylation of protein N termini, and oxidation of Met were considered as variable modifications. Trypsin as the proteolytic enzyme with up to 2 missed cleavages was allowed. The maximum false discovery rate was 0.01 and a minimum peptide length of 7 amino acids was required. All other parameters were default parameters of MaxQuant. Quantitative normalized ratios were calculated by MaxQuant and used for further data analysis.

### Lipidomics

In order to determine the change in lipid composition, Hek293T wt cells together with three different Hek293TΔSPP clones were analyzed by lipidomics. At day 0, cells were seeded onto PLL-coated 100 mm dishes in DMEM supplemented with 10% FBS. At day 1, cells were first washed two times with 1x PBS followed by sterol-depletion in medium containing DMEM supplemented with 5% lipoprotein deficient serum (LPDS) (see below). At day of harvest (day 2 after seeding), membrane fractions were prepared by subcellular fractionation (see below). The last centrifugation step was performed for 60 min at 21,000 x *g* at 4°C in a table-top centrifuge. Membranes were resuspended in methanol and subjected to lipid extractions using an acidic Bligh & Dyer, except from plasmalogens, which were extracted under neutral conditions as described in (50). Lipid standards were added prior to extractions, using a master mix containing phosphatidylcholine (13:0/13:0, 14:0/14:0, 20:0/20:0; 21:0/21:0; Avanti Polar Lipids, Alabaster, AL, USA), sphingomyelin (SM, d18:1 with N-acylated 13:0, 17:0, 25:0, semi-synthesized (50)), D_6_-cholesterol (Cambridge Isotope Laboratory), phosphatidylinositol (PI, 16:0/16:0 and 17:0/20:4; Avanti Polar Lipids, Alabaster, AL, USA), phosphatidylethanolamine (PE), phosphatidylserine (PS) and phosphatidylglycerol (PG) (14:1/14:1, 20:1/20:1, 22:1/22:1, semi-synthesized (50), diacylglycerol (DAG, 17:0/17:0, Larodan), cholesterol ester (CE, 9:0, 19:0, 24:1, Sigma-Aldrich, St. Louis, MO, USA), triacylglycerol (TAG, D_5_-mix, LM-6000/D_5_-17:0,17:1,17:1; Avanti Polar Lipids, Alabaster, AL, USA), ceramide (Cer) and glucosylceramide (HexCer) (both d18:1 with N-acylated 15:0, 17:0, 25:0, semi-synthesized (50), lactosylceramide (LacCer, d18:1 with N-acylated C12 fatty acid; Avanti Polar Lipids, Alabaster, AL, USA), phosphatidic acid (PA, 17:0/20:4; Avanti Polar Lipids, Alabaster, AL, USA), and lyso-phosphatidylcholine (LPC, 17:1; Avanti Polar Lipids, Alabaster, AL, USA). PE plasmalogen (PE P-)-containing standard mix was supplemented with PE P-mix 1 (16:0p/15:0, 16:0p/19:0, 16:0p/ 25:0), PE P-mix 2 (18:0p/15:0, 18:0p/19:0, 18:0p/25:0), and PE P-Mix 3 (18:1p/15:0, 18:1p/19:0, 18:1p/25:0). Semi-synthesis of PE P- was performed as described in (51). Lipid extracts were resuspended in 60 µl methanol and samples were analyzed on an AB SCIEX QTRAP 6500+ mass spectrometer (Sciex, Framingham, MA, USA) with chip-based (HD-D ESI Chip; Advion Biosciences, Ithaca, NY, USA) nano-electrospray infusion and ionization via a Triversa Nanomate (Advion Biosciences, Ithaca, NY, USA) as previously described (51). Resuspended lipid extracts were diluted 1:10 in 96-well plates (Eppendorf twin tec 96, colorless, Z651400-25A; Sigma-Aldrich, St. Louis, MO, USA) prior to measurement. Lipid classes were analyzed in positive ion mode applying either specific precursor ion (PC, lyso-PC, SM, cholesterol, Cer, HexCer, Hex2Cer, and PE-P) or neutral loss (PE, PS, PI, PG, and PA) scanning as described in (51). Data evaluation was performed using LipidView (RRID: SCR_017003; Sciex, Framingham, MA, USA) and an in-house-developed software (ShinyLipids).

### Quantitative real-time PCR

Quantitative real-time PCR was used to assess the SQS mRNA level (transcribed from the *FDFT1* gene) in Hek293T wt cells or cells deficient for SPP, TRC8, MARCH6, or TRC8/MARCH6-double deficient cells. Therefore, RNA was isolated from confluent cells from one well of a 6-well plate with the NucleoSpin RNA isolation kit (Macherey-Nagel) according to manufacturer’s protocol. 2 µg RNA were reverse transcribed using the RevertAid First Strand cDNA Synthesis Kit (Thermo Scientific). Quantitative PCR was performed using the SensiFAST SYBR No-ROX kit (Bioline) according to the manufacturer’s protocol and the LightCycler480 Instrument II (Roche) with the following cycle settings: pre-incubation for 2 min at 95°C, followed by 40 cycles amplification at 95°C for 5 sec, 57°C for 10 sec, and 72°C for 25 sec. Each reaction was performed in technical triplicate. TATAA-box binding protein (TBP) and β-2 microglobulin (β2M) were used as reference genes. Primers used for TBP: 5’-CCGGCTGTTTAACTTCGTT and 5’-ACGCCAAGAAACAGTGATGC; β2M: 5’-CACGTCATCCAGCAGAGAAT and 5’-TGCTGCTTACATGTCTCGAT; SQS/*FDFT1*: 5’-GAGGACTTCCCAACGATCTCC and 5’-AACTCTGCCATCCCAATGCC. Relative quantification of gene expression was calculated as described previously (52).

### Preparation of LPDS, cholesterol inclusion complexes and cholesterol-depletion and repletion assay

For the manipulation of the cellular cholesterol content, Hek293T cells were pre-treated and treated with DMEM containing 5% LPDS. LPDS was prepared from FBS by density gradient ultracentrifugation using solid potassium bromide according to (53) with minor modifications. The density adjusted FBS was subjected to ultracentrifugation for 25 h at 310,500 x *g* at 10°C in a 70Ti rotor (Beckman Coulter) before dialysis. The dialyzed fraction was sterile filtered and frozen at −20°C.

Cholesterol repletion was carried out by incubating cells in the presence of cholesterol/methyl-β-cyclodextrin inclusion complexes (Chol/CD) which were prepared with minor modifications according to (54). Cholesterol (Sigma-Aldrich) was dissolved in isopropanol:chloroform (2:1) and added in small aliquots to a stirring 5% (w/v) methyl-β-cyclodextrin (Sigma-Aldrich) solution in a water bath heated to 80°C. The solution was cooled down, sterile filtered and kept at 4°C protected from light until use. The cholesterol to methyl-β-cyclodextrin molar ratio was 10:1.

For experiments involving sterol depletion and repletion, cells were seeded onto PLL-coated plates. After 24 hours, cells were washed two times with 1x PBS before switching to sterol-depletion medium containing 5% LPDS, supplemented with 5 µM mevastatin and 50 µM mevalonate (Sigma-Aldrich). After overnight sterol depletion in this medium, cells were treated with fresh sterol-depletion medium either in absence or in presence of 30 µg/ml Chol/CD complexes or other test agents.

### Crystal violet cytotoxicity assay

For crystal violet cell viability assay, cells were seeded onto PLL-coated 24-well plates, sterol-depleted overnight with medium containing LPDS, mevastatin and mevalonate and treated with or without 30 µg/ml Chol/CD. At three hours after treatment, cells were washed with 1x PBS and fixed for 10 min at 4°C with cold methanol/acetone (1:1; 1 ml/well). Cells were stained with 0.1% crystal violet in 25% methanol for 15 min followed by washing four times with tap water. Afterwards, the stained cells were air dried overnight and representative images were acquired using a Zeiss Axio Observer Z1 widefield microscope equipped with a LD-Plan-NEOFLUAR 20x/0.4 Air Ph2 objective at 20x magnification. For quantitative measurement, adsorbed dye was solubilized with methanol containing 0.1% SDS by gentle agitation for two hours. Optical density at 570 nm (OD_570_) from three biological replicates and each replicate in technical duplicate was measured using the Infinite M1000 microplate reader (Tecan Trading AG, Switzerland) and accompanying software i-control (v.1.10.4.0). For quantification, the mean OD for each cell line and condition tested was corrected by subtracting the mean OD_570_ of blank wells. The OD_570_ of FBS-treated cells for each cell line was set to 1.0 and the respective fold-change under treatment conditions for each cell line was calculated.

### Cycloheximide chase experiments

Cycloheximide (100 µg/ml) chase was performed 24 h after transfection of Hek293T cells or after doxycycline induction of Hek293 Flp-In T-REx cells. For chase of endogenous proteins, cycloheximide was added 48 h after seeding of Hek293T cells. Following the addition of the compound in medium onto cells, lysates were prepared at indicated time points and protein abundance was analyzed by western blotting. Time point 0 was immediately harvested and did not receive cycloheximide.

### Subcellular fractionation

In order to isolate cellular membranes, subcellular fractionation was performed. Therefore, cells from 6-well or 100 mm dishes were washed with 1x PBS followed by adding 1 ml PBS-EDTA to detach cells. Cells were centrifuged for 10 min at 400 x *g* at 4°C, the supernatant was removed, and the cell pellet was resuspended in 1 ml hypotonic buffer supplemented with 10 µg/ml PMSF and protease inhibitor cocktail (Roche). Cells were incubated on ice for 10 min, lysed by passing 5 times through a 27-Gauge needle, followed by centrifugation for 5 min at 1,000 x *g* at 4°C to pellet cells and debris. To pellet membranes, the post-nuclear supernatant was centrifuged for 30 min at 100,000 x *g* at 4°C using a S120AT2 rotor (Beckman Coulter). The resulting pellet containing the membranes was resuspended in SDS sample buffer (see below).

Fractionation into plasma membrane and the ER-containing endomembrane fraction for the lipidome analysis was performed using the Minute™ Plasma Membrane Protein Isolation and Cell Fractionation Kit (Invent Biotechnologies, Inc.) according to manufacturer’s instructions.

### Fluorescence microscopy and fluid phase uptake assay

For immunofluorescence staining, cells were first carefully washed with 1x PBS followed by fixation with 0.5 ml/well 4% formaldehyde in PBS for 20 min with gentle agitation. Afterwards, cells were washed three times with 1x PBS and then permeabilized with 0.2 ml/well 0.5% Triton X-100 in PBS for 10 min at 4°C. Following washing with 1x PBS (three times), unspecific binding sites were blocked by incubating the cells with blocking buffer (20% FBS in 1x PBS) for 30 min. Primary antibody in blocking buffer supplemented with 0.01% Triton X-100 was applied for 1 h with gentle agitation followed by washing three times with 1x PBS, 5 min each. After this, the secondary antibody at respective dilution in blocking buffer was applied onto the cells and incubated for 1 h with agitation. From this step onwards, cells were protected from light to prevent antibody bleaching. Cells were washed three times with 1x PBS, 5 min each and mounted in Fluoromount-G (Southern Biotech). Confocal analysis was performed on a Zeiss LSM 780 inverted microscope. Lasers used were a 458 nm, 488 nm, 514 nm Argon laser (25 mW) and a 561 nm DPSS laser (20 mW). Images were acquired using a Plan-APOCHROMAT 63x/1.4 oil objective at 1024 x 1024 pixels with line average 4 and a pinhole size of 1 Airy unit using the ZEN 2012 (v. 1.1.2.0) software. For image stacks, series of 8-12 sections with 0.5 µm along the Z-axis were taken. Images were processed and maximum intensity projections created using Fiji.

In order to assess the influence of cholesterol on dextran internalization, fluid phase uptake assays were done. For this, cells were kept overnight in sterol depleted medium (see above). The next day, cells were either left under sterol depletion or in repletion medium with cholesterol for 3 hours. For the last 30 min, cells were incubated with 10 kDa dextran-Alexa568 and 70 kDa dextran-Oregon Green 488 at 0.1 mg/ml each. Subsequently, cells were washed two times with ice cold PBS and fixed with cold 4% formaldehyde in PBS for 15 min. After fixation, cells were washed in PBS, stained with 5 µg/ml Hoechst 33342 stain in PBS for 5 min, and finally rinsed with PBS prior to microscopy. Cellular uptake of fluorescently labelled dextran was analyzed by confocal microscopy using a Zeiss LSM 780 inverted microscope and quantified using Fiji/ImageJ (http://rsb.info.nih.gov/ij/) according to (55).

### Immunoprecipitation and TUBE-ubiquitin pulldown

For immunoprecipitation, cells were washed once with 1x PBS followed by treatment with the membrane-permeable crosslinker DSP (2 mM) in PBS for 30 min on ice. Tris-HCl, pH 7.4 was added to a final concentration of 20 mM for 15 min to quench the cross-linking reaction. Cells were washed with 1x PBS, harvested in PBS-EDTA followed by centrifugation for 3 min at 900 x *g* at 4°C. The supernatant was discarded, and the pellet was lysed in 1 ml solubilization buffer (50 mM HEPES-KOH, pH 7.4, 150 mM NaCl, 1.5 mM MgCl_2_, 1 mM EGTA, 10% glycerol) containing 10 µg/ml PMSF, complete protease inhibitor cocktail and 1% CHAPS. Cells were lysed for 30 min on ice followed by centrifugation for 10 min at 13,000 rpm at 4°C. The supernatant was diluted 1:1 with solubilization buffer without detergent and lysates were pre-cleared by incubation with Protein G sepharose beads (GE Healthcare) on a rotation wheel for 2 h, at 4°C. For immunoprecipitation, pre-cleared lysates were spun for 3 min at 4,000 x *g* at 4°C, transferred to a clean tube and incubated with monoclonal anti-HA agarose beads (Sigma-Aldrich) for 4 h, at 4°C. Beads were washed three times with wash buffer (50 mM HEPES-KOH, pH 7.4, 150 mM NaCl, 1.5 mM MgCl_2_, 1 mM EGTA, 10% glycerol) containing 0.1% CHAPS and eluted in 2x SDS-sample buffer (see below).

For determining the ubiquitination of endogenous SQS, Hek293T, Hek293TΔSPP and Hek293TΔTRC8 cells were seeded onto PLL-coated 100 mm cell culture dishes. Cells were washed two times with 1x PBS and treated overnight with medium containing 5% (v/v) LPDS. Prior to harvest, the proteasome inhibitor epoxomicin (final 5 µM) was added to each dish and one dish of each cell line received 30 µg/ml Chol/CD for 3 hours. Cells were washed once with 1x PBS and harvested in 1x PBS-EDTA supplemented with 20 mM *N*-ethylmaleimide (NEM) to block deubiquitinating enzymes followed by centrifugation for 3 min at 900 x *g* at 4°C. Cells were lysed for 20 min on ice in 500 µl solubilization buffer containing freshly added protease inhibitor cocktail, PMSF, 1% Triton X-100 and 20 mM NEM. Lysates were clarified by centrifugation for 15 min at full speed at 4°C and the supernatant was diluted 1:1 with solubilization buffer without detergent. Lysates were pre-cleared at 4°C for 1 h on a rotating wheel using 30 µl Protein-S agarose beads (Merck Millipore) slurry. Beads were centrifuged for 3 min at 4000 x *g* and the pre-cleared lysate transferred to a fresh tube. Affinity purification/enrichment of ubiquitinated species was performed using 20 µl TUBE2 agarose bead (LifeSensors) slurry, incubating overnight at 4°C on a rotating wheel. Beads were washed three times with TBS-T (50 mM Tris-HCl, pH 7.4, 150 mM NaCl, 0.1% Tween-20). Proteins were eluted by heating to 65°C for 15 min in 20 µl 1x SDS-sample buffer. Samples were subjected to SDS-PAGE and western blotting (see below).

### SDS-PAGE and western blotting

Proteins were solubilized in SDS sample buffer (50 mM Tris-HCl, pH 6.8, 10 mM EDTA, pH 8.0, 5% glycerol, 2% SDS, 0.01% bromophenol blue) containing 5% β-mercaptoethanol. Samples were heated to 65°C for 15 min with agitation, except for immunoblots of Hrd1 which were incubated at 37°C for 30 min. For western blotting, proteins were separated by reducing Tris-glycine SDS-PAGE and transferred to polyvinylidene difluoride (PVDF, Merck Millipore) membranes followed by enhanced chemiluminescence analysis (Advansta) using the LAS4000 system (Fuji). For reprobing, PVDF membranes were incubated in glycine stripping buffer (100 mM glycine, 20 mM Mg(OAc)_2_, 50 mM KCl, 1% Tween-20, 2.5% SDS), pH 2.2, for 20 min followed by washing 6 times for 10 min with TBS-T before processing at outlined above.

### Statistical analysis

Quantification of western blots was performed using ImageJ (http://rsb.info.nih.gov/ij/). Statistical analysis was performed using GraphPad Prism v.5.00 (GraphPad Software, Inc.). Differences between two means in steady state analysis was determined by two-tailed unpaired Student’s *t*-test. The difference between multiple means in cycloheximide chase assays was determined by two-way ANOVA test followed by Bonferroni *post hoc* test. If not stated otherwise, data are represented as mean ± SEM and result from at least three independent biological replicates. Significance levels were as follows: \**P* ≤ 0.05, \*\**P* < 0.01, \*\*\**P* < 0.001.

*This article contains supporting information*.

## Acknowledgments

We thank Bernhard Dobberstein and Robert M. Gemmill for reagents, Wilhelm Palm and Colin Adrain, Bernd Schröder and Torben Mentrup for advice and critical reading of the manuscript, Iris Leibrecht for technical support with the lipidome analysis and Thomas Ruppert (ZMBH MS facility) for MS/MS analysis.

## Author contribution

R.H., D.A. and M.K.L. conceptualized the project. R.H., D.A. and M.K.L. designed the research and wrote the manuscript. R.H., D.A. and M.C. performed the experiments and analyzed the data. C.L., D.K., and B.B. performed the lipidome analysis and analyzed the data. All authors contributed and approved the final version of the manuscript.

## Abbreviations

CHX: Cycloheximide
EMC: ER membrane protein complex
ER: Endoplasmic reticulum
ERAD: ER-associated degradation
FBS: fetal bovine serum
FPP: farnesyl pyrophosphate
HMGCR: HMG-CoA reductase
HO1: heme oxygenase 1
LPDS: lipoprotein deficient serum
SILAC: stable-isotope labeling by amino acids in cell culture
SQLE: squalene epoxisade
SQS: squalene synthase
SPP: signal peptide peptidase
SREBP: sterol regulatory element binding protein
SSD: sterol sensing domain
TM: transmembrane
WB: western blot

## References

1. Goldstein, J. L., and Brown, M. S. (1990) Regulation of the mevalonate pathway. Nature 343, 425–430

2. Edwards, P. A., and Ericsson, J. (1999) Sterols and isoprenoids: signaling molecules derived from the cholesterol biosynthetic pathway. Annu Rev Biochem 68, 157–185

3. Brown, M. S., Radhakrishnan, A., and Goldstein, J. L. (2018) Retrospective on Cholesterol Homeostasis: The Central Role of Scap. Annu Rev Biochem 87, 783–807

4. Schumacher, M. M., and DeBose-Boyd, R. A. (2021) Posttranslational Regulation of HMG CoA Reductase, the Rate-Limiting Enzyme in Synthesis of Cholesterol. Annu Rev Biochem 90, 659–679

5. Gill, S., Stevenson, J., Kristiana, I., and Brown, A. J. (2011) Cholesterol-dependent degradation of squalene monooxygenase, a control point in cholesterol synthesis beyond HMG-CoA reductase. Cell Metab 13, 260–273

6. Foresti, O., Ruggiano, A., Hannibal-Bach, H. K., Ejsing, C. S., and Carvalho, P. (2013) Sterol homeostasis requires regulated degradation of squalene monooxygenase by the ubiquitin ligase Doa10/Teb4. Elife 2, e00953

7. Zelcer, N., Sharpe, L. J., Loregger, A., Kristiana, I., Cook, E. C., Phan, L., Stevenson, J., and Brown, A. J. (2014) The E3 ubiquitin ligase MARCH6 degrades squalene monooxygenase and affects 3-hydroxy-3-methyl-glutaryl coenzyme A reductase and the cholesterol synthesis pathway. Mol Cell Biol 34, 1262–1270

8. Lemberg, M. K., and Strisovsky, K. (2021) Maintenance of organellar protein homeostasis by ER-associated degradation and related mechanisms. Mol Cell 81, 2507–2519

9. Coates, H. W., Capell-Hattam, I. M., and Brown, A. J. (2021) The mammalian cholesterol synthesis enzyme squalene monooxygenase is proteasomally truncated to a constitutively active form. J Biol Chem 296, 100731

10. Howe, V., Sharpe, L. J., Alexopoulos, S. J., Kunze, S. V., Chua, N. K., Li, D., and Brown, A. J. (2016) Cholesterol homeostasis: How do cells sense sterol excess? Chem Phys Lipids 199, 170–178

11. Rawson, R. B., Zelenski, N. G., Nijhawan, D., Ye, J., Sakai, J., Hasan, M. T., Chang, T. Y., Brown, M. S., and Goldstein, J. L. (1997) Complementation cloning of S2P, a gene encoding a putative metalloprotease required for intramembrane cleavage of SREBPs. Mol. Cell 1, 47–57

12. Gardner, R. G., Shearer, A. G., and Hampton, R. Y. (2001) In vivo action of the HRD ubiquitin ligase complex: mechanisms of endoplasmic reticulum quality control and sterol regulation. Mol Cell Biol 21, 4276–4291

13. Jo, Y., Lee, P. C., Sguigna, P. V., and DeBose-Boyd, R. A. (2011) Sterol-induced degradation of HMG CoA reductase depends on interplay of two Insigs and two ubiquitin ligases, gp78 and Trc8. Proceedings of the National Academy of Sciences of the United States of America 108, 20503–20508

14. Jiang, L. Y., Jiang, W., Tian, N., Xiong, Y. N., Liu, J., Wei, J., Wu, K. Y., Luo, J., Shi, X. J., and Song, B. L. (2018) Ring finger protein 145 (RNF145) is a ubiquitin ligase for sterol-induced degradation of HMG-CoA reductase. J Biol Chem 293, 4047–4055

15. Menzies, S. A., Volkmar, N., van den Boomen, D. J., Timms, R. T., Dickson, A. S., Nathan, J. A., and Lehner, P. J. (2018) The sterol-responsive RNF145 E3 ubiquitin ligase mediates the degradation of HMG-CoA reductase together with gp78 and Hrd1. Elife 7

16. Avci, D., and Lemberg, M. K. (2015) Clipping or Extracting: Two Ways to Membrane Protein Degradation. Trends Cell Biol 25, 611–622

17. Weihofen, A., Binns, K., Lemberg, M. K., Ashman, K., and Martoglio, B. (2002) Identification of signal peptide peptidase, a presenilin-type aspartic protease. Science 296, 2215–2218.

18. Chen, C., Malchus, N. S., Hehn, B., Stelzer, W., Avci, D., Langosch, D., and Lemberg, M. K. (2014) Signal peptide peptidase functions in ERAD to cleave the unfolded protein response regulator XBP1u. EMBO J 33, 2492–2506

19. Boname, J. M., Bloor, S., Wandel, M. P., Nathan, J. A., Antrobus, R., Dingwell, K. S., Thurston, T. L., Smith, D. L., Smith, J. C., Randow, F., and Lehner, P. J. (2014) Cleavage by signal peptide peptidase is required for the degradation of selected tail-anchored proteins. J Cell Biol 205, 847–862

20. Stagg, H. R., Thomas, M., van den Boomen, D., Wiertz, E. J., Drabkin, H. A., Gemmill, R. M., and Lehner, P. J. (2009) The TRC8 E3 ligase ubiquitinates MHC class I molecules before dislocation from the ER. J Cell Biol 186, 685–692

21. Stefanovic-Barrett, S., Dickson, A. S., Burr, S. P., Williamson, J. C., Lobb, I. T., van den Boomen, D. J., Lehner, P. J., and Nathan, J. A. (2018) MARCH6 and TRC8 facilitate the quality control of cytosolic and tail-anchored proteins. EMBO Rep 19

22. Hsu, F. F., Yeh, C. T., Sun, Y. J., Chiang, M. T., Lan, W. M., Li, F. A., Lee, W. H., and Chau, L. Y. (2015) Signal peptide peptidase-mediated nuclear localization of heme oxygenase-1 promotes cancer cell proliferation and invasion independent of its enzymatic activity. Oncogene 34, 2360–2370

23. Avci, D., Malchus, N. S., Heidasch, R., Lorenz, H., Richter, K., Nessling, M., and Lemberg, M. K. (2019) The intramembrane protease SPP impacts morphology of the endoplasmic reticulum by triggering degradation of morphogenic proteins. J Biol Chem 294, 2786–2800

24. Yucel, S. S., Stelzer, W., Lorenzoni, A., Wozny, M., Langosch, D., and Lemberg, M. K. (2019) The Metastable XBP1u Transmembrane Domain Defines Determinants for Intramembrane Proteolysis by Signal Peptide Peptidase. Cell reports 26, 3087–3099 e3011

25. Guna, A., Volkmar, N., Christianson, J. C., and Hegde, R. S. (2018) The ER membrane protein complex is a transmembrane domain insertase. Science 359, 470–473

26. Volkmar, N., Thezenas, M. L., Louie, S. M., Juszkiewicz, S., Nomura, D. K., Hegde, R. S., Kessler, B. M., and Christianson, J. C. (2019) The ER membrane protein complex promotes biogenesis of sterol-related enzymes maintaining cholesterol homeostasis. J Cell Sci 132

27. Aizawa, S., Okamoto, T., Sugiyama, Y., Kouwaki, T., Ito, A., Suzuki, T., Ono, C., Fukuhara, T., Yamamoto, M., Okochi, M., Hiraga, N., Imamura, M., Chayama, K., Suzuki, R., Shoji, I., Moriishi, K., Moriya, K., Koike, K., and Matsuura, Y. (2016) TRC8-dependent degradation of hepatitis C virus immature core protein regulates viral propagation and pathogenesis. Nat Commun 7, 11379

28. Brown, M. S., and Goldstein, J. L. (1980) Multivalent feedback regulation of HMG CoA reductase, a control mechanism coordinating isoprenoid synthesis and cell growth. J Lipid Res 21, 505–517

29. Nakanishi, M., Goldstein, J. L., and Brown, M. S. (1988) Multivalent control of 3-hydroxy-3-methylglutaryl coenzyme A reductase. Mevalonate-derived product inhibits translation of mRNA and accelerates degradation of enzyme. J Biol Chem 263, 8929–8937

30. Sever, N., Song, B. L., Yabe, D., Goldstein, J. L., Brown, M. S., and DeBose-Boyd, R. A. (2003) Insig-dependent ubiquitination and degradation of mammalian 3-hydroxy-3-methylglutaryl-CoA reductase stimulated by sterols and geranylgeraniol. J Biol Chem 278, 52479–52490

31. Irisawa, M., Inoue, J., Ozawa, N., Mori, K., and Sato, R. (2009) The sterol-sensing endoplasmic reticulum (ER) membrane protein TRC8 hampers ER to Golgi transport of sterol regulatory element-binding protein-2 (SREBP-2)/SREBP cleavage-activated protein and reduces SREBP-2 cleavage. J Biol Chem 284, 28995–29004

32. Lee, J. P., Brauweiler, A., Rudolph, M., Hooper, J. E., Drabkin, H. A., and Gemmill, R. M. (2010) The TRC8 ubiquitin ligase is sterol regulated and interacts with lipid and protein biosynthetic pathways. Mol Cancer Res 8, 93–106

33. Sandvig, K., Torgersen, M. L., Raa, H. A., and van Deurs, B. (2008) Clathrin-independent endocytosis: from nonexisting to an extreme degree of complexity. Histochem Cell Biol 129, 267–276

34. Skotland, T., and Sandvig, K. (2019) The role of PS 18:0/18:1 in membrane function. Nat Commun 10, 2752

35. Wei, J. W., Cai, J. Q., Fang, C., Tan, Y. L., Huang, K., Yang, C., Chen, Q., Jiang, C. L., and Kang, C. S. (2017) Signal Peptide Peptidase, Encoded by HM13, Contributes to Tumor Progression by Affecting EGFRvIII Secretion Profiles in Glioblastoma. CNS Neurosci Ther 23, 257–265

36. Hsu, F. F., Chou, Y. T., Chiang, M. T., Li, F. A., Yeh, C. T., Lee, W. H., and Chau, L. Y. (2018) Signal peptide peptidase promotes tumor progression via facilitating FKBP8 degradation. Oncogene

37. Hua, X., Sakai, J., Brown, M. S., and Goldstein, J. L. (1996) Regulated cleavage of sterol regulatory element binding proteins requires sequences on both sides of the endoplasmic reticulum membrane. J Biol Chem 271, 10379–10384.

38. Wahrle, S., Das, P., Nyborg, A. C., McLendon, C., Shoji, M., Kawarabayashi, T., Younkin, L. H., Younkin, S. G., and Golde, T. E. (2002) Cholesterol-dependent gamma-secretase activity in buoyant cholesterol-rich membrane microdomains. Neurobiol Dis 9, 11–23

39. Yang, G., Zhou, R., Zhou, Q., Guo, X., Yan, C., Ke, M., Lei, J., and Shi, Y. (2019) Structural basis of Notch recognition by human gamma-secretase. Nature 565, 192–197

40. Kaymak, I., Maier, C. R., Schmitz, W., Campbell, A. D., Dankworth, B., Ade, C. P., Walz, S., Paauwe, M., Kalogirou, C., Marouf, H., Rosenfeldt, M. T., Gay, D. M., McGregor, G. H., Sansom, O. J., and Schulze, A. (2020) Mevalonate Pathway Provides Ubiquinone to Maintain Pyrimidine Synthesis and Survival in p53-Deficient Cancer Cells Exposed to Metabolic Stress. Cancer Res 80, 189–203

41. Brauweiler, A., Lorick, K. L., Lee, J. P., Tsai, Y. C., Chan, D., Weissman, A. M., Drabkin, H. A., and Gemmill, R. M. (2007) RING-dependent tumor suppression and G2/M arrest induced by the TRC8 hereditary kidney cancer gene. Oncogene 26, 2263–2271

42. Snapp, E. L., Sharma, A., Lippincott-Schwartz, J., and Hegde, R. S. (2006) Monitoring chaperone engagement of substrates in the endoplasmic reticulum of live cells. Proceedings of the National Academy of Sciences of the United States of America 103, 6536–6541

43. Niemeyer, J., Mentrup, T., Heidasch, R., Muller, S. A., Biswas, U., Meyer, R., Papadopoulou, A. A., Dederer, V., Haug-Kroper, M., Adamski, V., Lullmann-Rauch, R., Bergmann, M., Mayerhofer, A., Saftig, P., Wennemuth, G., Jessberger, R., Fluhrer, R., Lichtenthaler, S. F., Lemberg, M. K., and Schroder, B. (2019) The intramembrane protease SPPL2c promotes male germ cell development by cleaving phospholamban. EMBO Rep 20

44. Durocher, Y., Perret, S., and Kamen, A. (2002) High-level and high-throughput recombinant protein production by transient transfection of suspension-growing human 293-EBNA1 cells. Nucleic Acids Res 30, E9

45. Heigwer, F., Kerr, G., and Boutros, M. (2014) E-CRISP: fast CRISPR target site identification. Nature methods 11, 122–123

46. Brinkman, E. K., Chen, T., Amendola, M., and van Steensel, B. (2014) Easy quantitative assessment of genome editing by sequence trace decomposition. Nucleic Acids Res 42, e168

47. Dehairs, J., Talebi, A., Cherifi, Y., and Swinnen, J. V. (2016) CRISP-ID: decoding CRISPR mediated indels by Sanger sequencing. Scientific reports 6, 28973

48. Conant, D., Hsiau, T., Rossi, N., Oki, J., Maures, T., Waite, K., Yang, J., Joshi, S., Kelso, R., Holden, K., Enzmann, B. L., and Stoner, R. (2022) Inference of CRISPR Edits from Sanger Trace Data. CRISPR J 5, 123–130

49. Bock, J., Kuhnle, N., Knopf, J. D., Landscheidt, N., Lee, J. G., Ye, Y., and Lemberg, M. K. (2022) Rhomboid protease RHBDL4 promotes retrotranslocation of aggregation-prone proteins for degradation. Cell reports 40, 111175

50. Ozbalci, C., Sachsenheimer, T., and Brugger, B. (2013) Quantitative analysis of cellular lipids by nano-electrospray ionization mass spectrometry. Methods Mol Biol 1033, 3–20

51. Paltauf, F., and Hermetter, A. (1994) Strategies for the synthesis of glycerophospholipids. Prog Lipid Res 33, 239–328

52. Pfaffl, M. W. (2001) A new mathematical model for relative quantification in real-time RT-PCR. Nucleic Acids Res 29, e45

53. Goldstein, J. L., Basu, S. K., and Brown, M. S. (1983) Receptor-mediated endocytosis of low-density lipoprotein in cultured cells. Methods Enzymol 98, 241–260

54. Klein, U., Gimpl, G., and Fahrenholz, F. (1995) Alteration of the myometrial plasma membrane cholesterol content with beta-cyclodextrin modulates the binding affinity of the oxytocin receptor. Biochemistry 34, 13784–13793

55. Palm, W., Araki, J., King, B., DeMatteo, R. G., and Thompson, C. B. (2017) Critical role for PI3-kinase in regulating the use of proteins as an amino acid source. Proceedings of the National Academy of Sciences of the United States of America 114, E8628–E8636

56. Cantagrel, V., and Lefeber, D. J. (2011) From glycosylation disorders to dolichol biosynthesis defects: a new class of metabolic diseases. J Inherit Metab Dis 34, 859–867

57. Lemberg, M. K., and Martoglio, B. (2002) Requirements for signal peptide peptidase-catalyzed intramembrane proteolysis. Mol. Cell 10, 735–744

