## Supplemental information and figures for "Intramembrane protease SPP defines a cholesterol-regulated abundance control of the mevalonate pathway enzyme SQS"

Supporting information includes supplementary figures 1-4 and supplementary file 1 (as a separate excel file).

**Supplementary file 1**

Complete list of proteins identified in high-salt washed and carbonate-extracted microsomes by SILAC-based quantitative proteomics. Lists for all ER-resident and all identified TA-proteins are given. Proteins enriched >20% are highlighted in blue; protein depleted >20% are highlighted yellow.

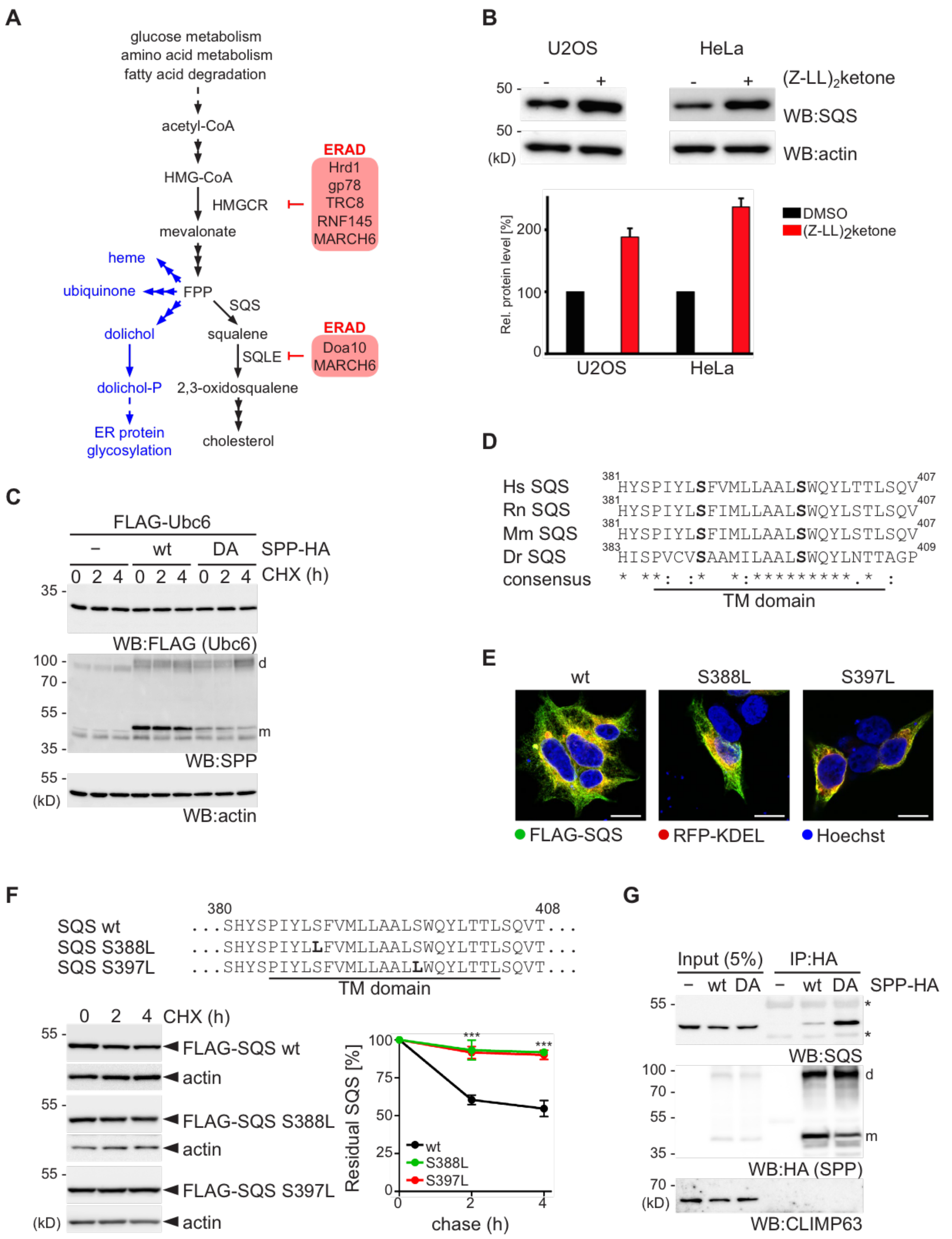


**Figure S1. TM region of SQS targets it for SPP-mediated ERAD**

**(A)** Simplified view of the branched mevalonate pathway leading to cholesterol biosynthesis by multiple steps (black arrows) and other isoprenoids including dolichol, which in its phosphorylated form (dolichol-P) serves as the lipid carrier for glycans in the ER (56), ubiquinone and heme (blue arrows) (1). Biosynthesis starts from acetyl-CoA, which derives from glucose metabolism and the breakdown of amino acids and fatty acids. The conversion of HMG-CoA to mevalonate by HMGCR is considered as the rate-limiting step. The intermediate FPP (farnesyl-pyrophosphate) can be shunted either into the non-sterol branch or the sterol-branch ultimately leading to cholesterol. The conversion of FPP to squalene by SQS is the first committed step in the sterol-branch. The over-all rate-limiting enzyme HMGCR is post-translationally regulated by ERAD involving multiple E3 ubiquitin ligases (red box) (7,12-15); SQLE and the yeast orthologue Erg1 are degraded by MARCH6/Doa10 (5-7). **(B)** Western blot (WB) showing steady state analysis of endogenous SQS in U2OS and HeLa cells upon 16 h treatment with $(Z\text{-}LL)_2$-ketone or DMSO. Actin was used as loading control. Quantification shows Mean ± SEM, *n*=3. **(C)** CHX chase of co-expressed FLAG-tagged Ubc6 with HA-tagged wt or DA-mutant SPP. Degradation of Ubc6 is not affected by SPP. Actin was used as loading control. **(D)** Sequence alignment of human (*Homo sapiens*, Hs), rat (*Rattus norvegicus*, Rn), mouse (*Mus musculus*, Mm), and zebrafish (*Danio rerio*, Dr) SQS homologs shows conserved serine residues (bold) in the TM domain. **(E)** Immunofluorescence microscopy of transiently transfected Hek293T cells with FLAG-tagged SQS constructs as indicated (green) co-expressing the luminal ER marker RFP-KDEL (red) showing ER localization. Five frames, each from Z-stacks with step sizes of 0.5 µm, are displayed. Hoechst 33342, nuclear counterstaining, is shown in blue. Scale bar, 10 µm. **(F)** Stability of ectopically expressed FLAG-tagged SQS wild-type (wt) and the TM domain mutants S388L and S397L was analyzed by CHX chase. Mean ± SEM, *n=3*. **(G)** Co-immunoprecipitation assay with ectopically expressed HA-tagged SPP wt or DA mutant and endogenous SQS. Asterisks represent antibody heavy and light chain.

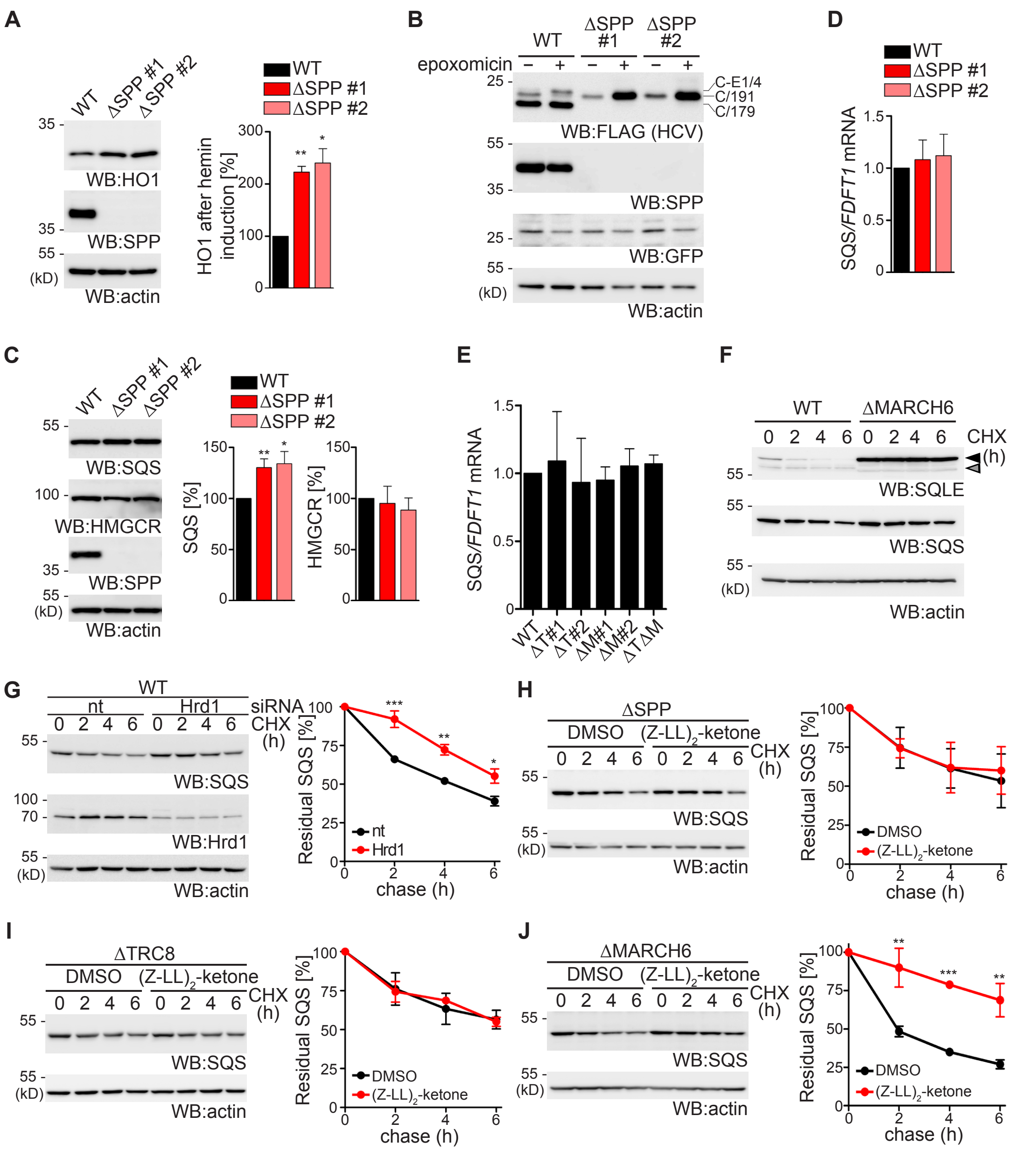


**Figure S2. SQS is degraded by a consorted action of SPP and TRC8.**

**(A)** Functional validation Hek293TΔSPP cells by analysis of steady state levels of endogenous HO1. Hek293T wt and two Hek293TΔSPP clones (#1, #2) were overnight treated with 20 µM hemin in order to induce expression of endogenous HO1 and subjected to immunoblotting. Endogenous HO1 is significantly accumulating in both ΔSPP clones. Immunoblot for SPP shows loss of protein. Only the monomeric form of SPP in Hek293T wt cells is shown. Mean ± SEM; *n=3*. **(B)** Functional validation of ΔSPP cells in respect to processing of the hepatitis C virus (HCV) nucleocapsid (core) protein. Hek293T wt and two Hek293TΔSPP clones (#1, #2) were transiently transfected with an N-terminally FLAG-tagged construct comprising the HCV core protein (C) followed by 4 amino acids of the E1 glycoprotein (E1/4) as has been used before (57). Cells were treated overnight in absence or presence of the proteasome inhibitor epoxomicin and subjected to immunoblotting. FLAG-tagged CE1/4 is first processed to the 191-residue immature core protein (C/191) by signal peptidase and then to 179 amino acid long mature core protein (C/179) by SPP. Cleavage of C/191 is abolished in ΔSPP cells leading to its degradation by ERAD as has been shown (27). Immunoblot for SPP shows loss of protein in both ΔSPP clones and the monomeric form of SPP in Hek293T wt cells is indicated. GFP was used as transfection control and actin was used as loading control. **(C)** Western blot (WB) showing the significant accumulation of endogenous SQS in two different Hek293TΔSPP clones (#1, #2) whereas HMGCR was unaffected. Only the monomeric from of SPP is shown. Actin was used as loading control. Mean ± SEM, *n=3*. **(D)** Quantitative real-time (qRT)-PCR for SQS (transcribed from the *FDFT1* gene) was performed on mRNA isolated from Hek293T wt and two Hek293T ΔSPP clones (#1, #2). Level of *SQS* mRNA were normalized to TBP and β2M and is shown as fold-change relative to wt mRNA level. Technical triplicates from three independent biological replicates were performed. Mean ± SEM, *n=3*. **(E)** qRT-PCR for SQS/*FDFT1* as in (**D**) but in two ΔTRC8 clones (ΔT#1, ΔT#2), two ΔMARCH6 clones (ΔM#1, ΔM#2), and one ΔTRC8ΔMARCH6 (ΔTΔM) double-knockout. Mean ± SEM, *n=3*. **(F)** CHX chase of endogenous SQLE and SQS in Hek293T wt and ΔMARCH6 cells showing striking stabilization of SQLE, but not of SQS. Black arrow; full length SQLE, grey arrow; SQLE-S. **(G)** CHX chase of endogenous SQS for indicated time-points in Hek293T wt cells transfected with non-targeting (nt) or Hrd1-specific siRNA. Mean ± SEM, *n=3*. **(H-J)** CHX chase of endogenous SQS in ΔSPP (**H**), ΔTRC8 (**I**), and ΔMARCH6 (**J**) Hek293T cells in presence of $(Z\text{-}LL)_2$-ketone or DMSO (vehicle). Mean ± SEM, *n=3*.

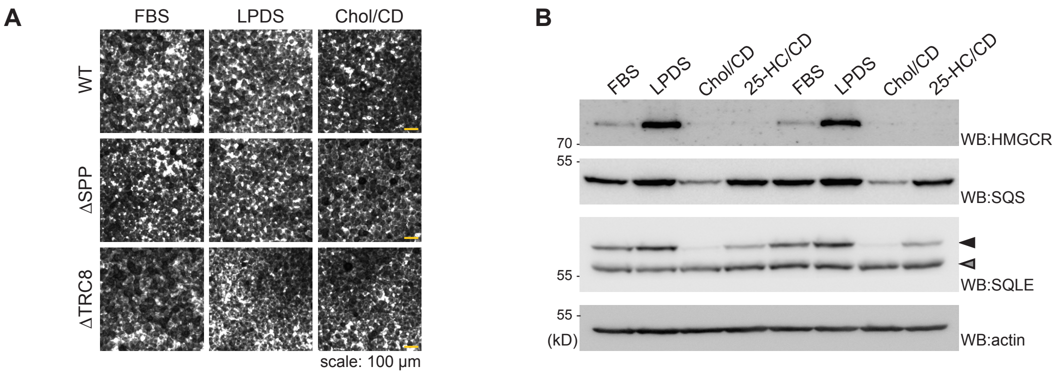


**Figure S3. Cholesterol but not 25-hydroxycholesterol triggers SPP-TRC8-dependent ERAD of SQS.**

**(A)** Crystal violet cytotoxicity assay shows that cell viability was not affected by the cholesterol depletion protocol outlined in Figure 3A. Hek293T wt, ΔSPP, or ΔTRC8 cells were seeded in 24-well plates, treated as indicated and assayed for cytotoxicity by crystal violet staining. **(B)** Western blot analysis showing effect of cholesterol and 25-hydroxycholesterol (25-HC) on the degradation of endogenous HMGCR, SQS and SQLE. Hek293T wt were grown in medium containing FBS or LPDS overnight and sterol depleted cells were treated for 5 h with either 30 µg/ml Chol/CD or 1 µg/ml 25-HC, as indicated. Western blot analysis of two independent experiments are shown. Black arrow; full length SQLE, grey arrow; SQLE-S.

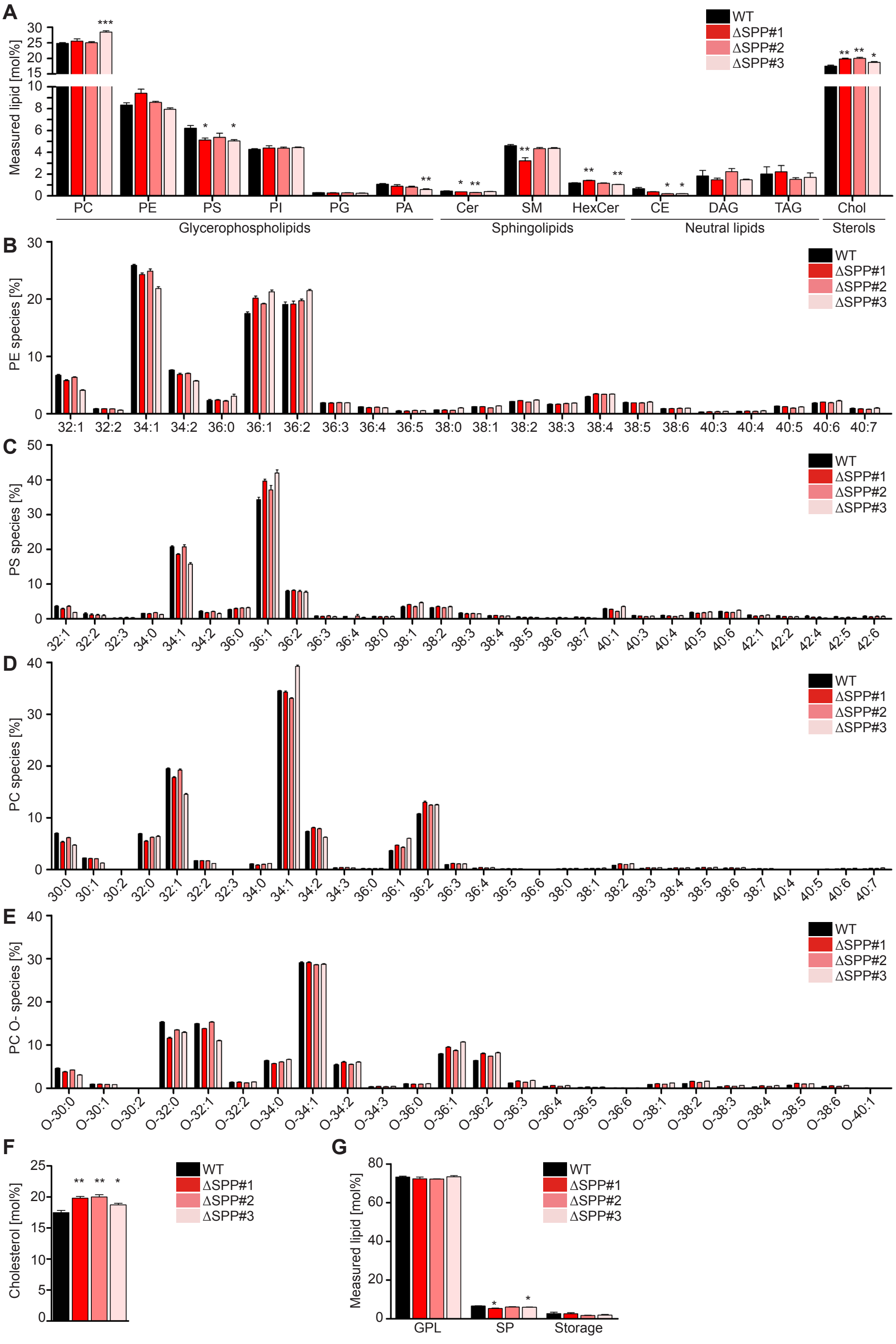


**Figure S4. Lipidomic profile of ΔSPP cells.**

**(A)** Lipid class profile of Hek293T wt and three Hek293TΔSPP (#1, #2, #3) cell lines. Lipid classes are standardized to all lipids measured and lipid categories are indicated. PC, phosphatidylcholine; PE, phosphatidylethanolamine; PS, phosphatidylserine; PI, phosphatidylinositol; PG, phosphatidylglycerol; PA, phosphatidic acid; Cer, ceramide; SM, sphingomyelin; HexCer, hexosylceramide; CE, cholesteryl ester; DAG, diacylglycerol; TAG, triacylglycerol. Mean ± SEM, *n*=4. **(B-E)** Molecular species profiles of PE (**B**), PS (**C**), PC (**D**) and PC O (E) are shown as % distributions within each class. Mean ± SEM; *n*=4. **(F)** Lipidomic analysis of total cholesterol levels in Hek293T wt and three independent ΔSPP clones (#1, #2, #3). Mean ± SEM, *n=4*. **(G)** Lipidomic analysis of functional lipid categories compared in Hek293T wt and three ΔSPP clones (#1, #2, #3). GPL, glycerophospholipids; SP, sphingolipids. Storage lipids include cholesteryl ester, diacylglycerol and triacylglycerol. Mean ± SEM, *n* = 4.
